# Pooled CRISPR screening at high sensitivity with an empirically designed sgRNA library

**DOI:** 10.1101/2020.04.25.061135

**Authors:** Luisa Henkel, Benedikt Rauscher, Barbara Schmitt, Jan Winter, Michael Boutros

## Abstract

Given their broad utility in functionally annotating genomes, the experimental design of genome-scale CRISPR screens can vary greatly and criteria for optimal experimental implementation and library composition are still emerging. In this study, we report advantages of conducting viability screens in selected Cas9 single cell clones in contrast to Cas9 bulk populations. We further systematically analyzed published CRISPR screens in human cells to identify single-guide (sg)RNAs with consistent high on-target and low off-target activity. Selected guides were collected in a new genome-scale sgRNA library, which efficiently identifies core and context-dependent essential genes. In summary, we show how empirically designed libraries in combination with an optimised experimental design increase the dynamic range in gene essentiality screens at reduced library coverage.

## Background

Over the past decades, genetic screens have been used extensively to interrogate gene function in an unbiased manner (Grimm 2004; Boutros and Ahringer 2008; Doench 2018). In recent years, CRISPR/Cas9 (Jinek et al. 2012; Mali et al. 2013) has emerged as a scalable method to introduce targeted gene knockouts with unprecedented efficiency and specificity. Genetic screens with the CRISPR/Cas9 system are now applied to probe the protein-coding and non-coding genomes of hundreds of cell types in different organisms (Wang et al. 2014; Shalem et al. 2014; Hart et al. 2015; Chen et al. 2015; Aguirre et al. 2016; John Liu et al. 2017). These experiments have led to new insights into many biological processes. Progress has been made especially in the field of cancer genetics, where genome-wide CRISPR screens have resulted in gene essentiality maps for hundreds of tumor cell lines (Meyers et al. 2017; Behan et al. 2019).

Despite these advances, currently no generally accepted design principles for sgRNA libraries (Sanson et al. 2018) and large-scale CRISPR screens exist and published experiments vary substantially in design and performance. This is particularly important as we reason that screens conducted with high CRISPR editing efficiency based on high Cas9 and sgRNA functionality allow screens at reduced coverage, which substantially lowers experimental efforts and costs. Furthermore, higher knockout efficiency is likely to increase the dynamic range of CRISPR screens, which in turn further allows to separate weak signals from screening noise and thus hit calling. In recent years, the design of sgRNAs and CRISPR libraries was improved based on design rules derived from comparing the nucleotide composition of active and non-active guides (Chen et al. 2018; Tzelepis et al. 2016; Hart et al. 2017), training models to identify predictive features of efficiently editing sgRNAs (Doench et al. 2014, 2016) and more recently also deep learning algorithms (Wang et al. 2019). While these efforts have greatly improved library performance, we reasoned that sequence predictions might not always fully reflect knockout performance and that many factors that influence sgRNA efficacy likely remain unknown and have thus not been considered in previous library designs. Potential further parameters might be DNA accessibility (Isaac et al. 2016; Horlbeck et al. 2016; Daer et al. 2017) or the presence of functional protein domains (Shi et al. 2015). Therefore, we consider using consistent and strong viability phenotypes of sgRNAs previously used in a diverse range of cell lines to be a powerful predictor of their functionality.

In this study, we generated a new library, termed the Heidelberg CRISPR library as a new tool for pooled genome-wide CRISPR/Cas9 screens. For its design, we selected guides with consistent high on-target and low off-target activity based on phenotypes in previously published CRISPR screens as recorded in the GenomeCRISPR database (Rauscher et al. 2017a). We show that empirical selection of sgRNAs also prioritizes guides with high sequence scores according to different design rules. In a next step, we show that our library efficiently identifies core and context-dependent essential genes by screening for essential genes in a cell line that was not considered for library design, the human HAP1 cell line. Screening in Cas9 single cell clones increased depletion phenotypes of essential genes compared to a Cas9 bulk population, increasing the overall dynamic range. Interestingly, the heterogeneity of editing efficiency of single cells in Cas9 bulk populations seems to more strongly interfere with editing efficiency compared to ploidy, since editing in haploid and diploid HAP1 cells was in a similar range. Furthermore, screening in selected single cell clones allowed hit calling at reduced library coverage, while essential genes were overall similar in all cell populations.

We believe that empirically designed libraries will be a useful extension to the current CRISPR/Cas9 toolkit. Our results further suggest that clonal cell populations with high Cas9 activity are attractive models for CRISPR/Cas9 screens at minimal library coverage, especially when no prior information is available to inform hit calling.

## RESULTS

### Data mining of sgRNA associated phenotypes allows empirical sgRNA design

We hypothesized that the very large number of results from previously published CRISPR screens could be utilized to systematically identify sgRNAs with high activity. Specifically, we reasoned that sgRNAs with strong and consistent effects across multiple experiments would intrinsically combine all known and unknown characteristics that guarantee high on-target activity. In addition, we assumed that we could avoid sgRNAs with off-target activity by comparing their phenotypes to other sgRNAs targeting the same gene. To this end we analyzed 439 genome-scale fitness screens (negative selection for viability) from GenomeCRISPR, a database that contains sgRNA phenotypes from CRISPR screens in human cells (Rauscher et al. 2017a, 2017b; Wang et al. 2014, 2015, 2017; Shalem et al. 2014; Aguirre et al. 2016; Munoz et al. 2016; Meyers et al. 2017; Tzelepis et al. 2016; Hart et al. 2015; Steinhart et al. 2017). We excluded all sgRNAs that could not be mapped to a protein coding transcript region of the latest (GRCh38.p10) human reference genome (Cunningham et al. 2019). In addition to the sgRNA phenotypes, we annotated each guide sequence with additional information including the number of targeted transcripts, the number of times the sgRNA was screened, the number of predicted off-targets (see Methods) and their GC content.

We next aimed to identify sgRNAs with high on-target activity (Figure 1). To this end we reanalysed each of the selected 439 screens in GenomeCRISPR using the BAGEL software (Hart and Moffat 2016). BAGEL uses reference sets of core essential and nonessential genes (Hart et al. 2017) to compute Bayes Factors that indicates whether a gene is more likely to be essential or nonessential. To exclude low-quality screens from further analysis, we generated precision-recall-curves for each screen to quantify how well core and nonessential reference genes could be separated based on the BAGEL-derived Bayes Factors (Hart et al. 2014). We retained all screens for which the area under the precision recall curve (AUC) was greater than 0.9 (406 out of 439; Supplementary Figures 1A-B).

**Figure 1:**
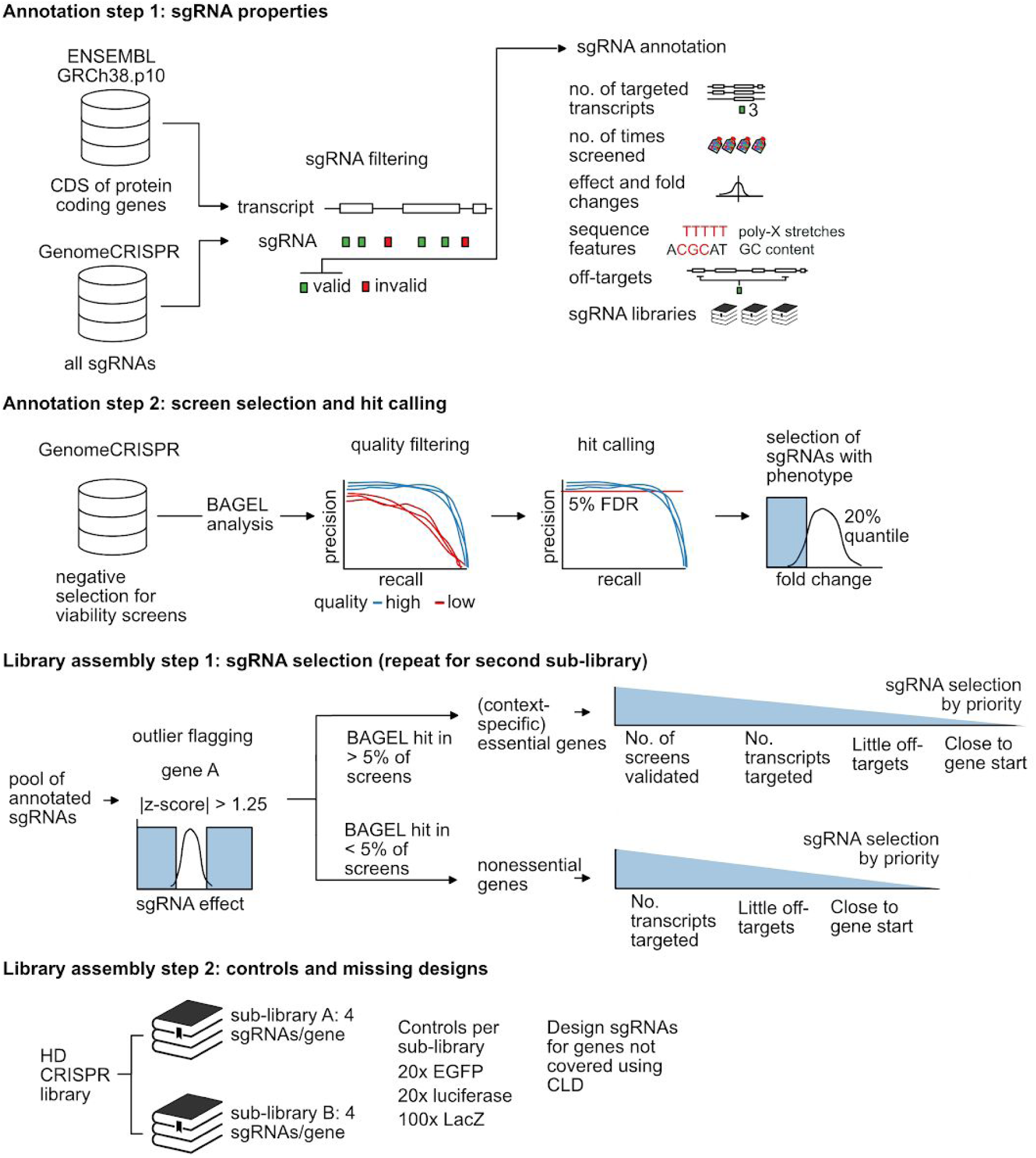
Empirical design of the HD CRISPR Library. Schematic illustration of the HD CRISPR Library design process. sgRNA sequences that were previously used in negative selection screens contained in the GenomeCRISPR database were annotated with information on rationally selected sequence and phenotype features. All negative selection screens were re-analyzed with BAGEL. High-quality experiments were selected based on how well reference core and nonessential gene sets could be separated in those screens. sgRNAs with high on-target activity were then determined as sequences that both target an essential gene (as determined by BAGEL) and that rank among the 20% most strongly depleted sequences in the screen. Next, sgRNAs that showed unexpected phenotypes compared to other sgRNAs targeting the same gene were flagged as outliers with potential off-target effects. sgRNA sequences were then selected from the resulting pool of sequences to design a genome-wide library consisting of two mutually exclusive sub-libraries A and B, prioritizing sgRNAs with high on-target and low off-target activity.

We then determined essential genes in each of these screens at 5% false discovery rate (FDR). We labelled each sgRNA as active if its target gene was essential according to BAGEL and if the sgRNA was among the 20% sgRNAs with the strongest fitness phenotypes in a screen (Figure 1; Figure 2 A-B).

**Figure 2:**
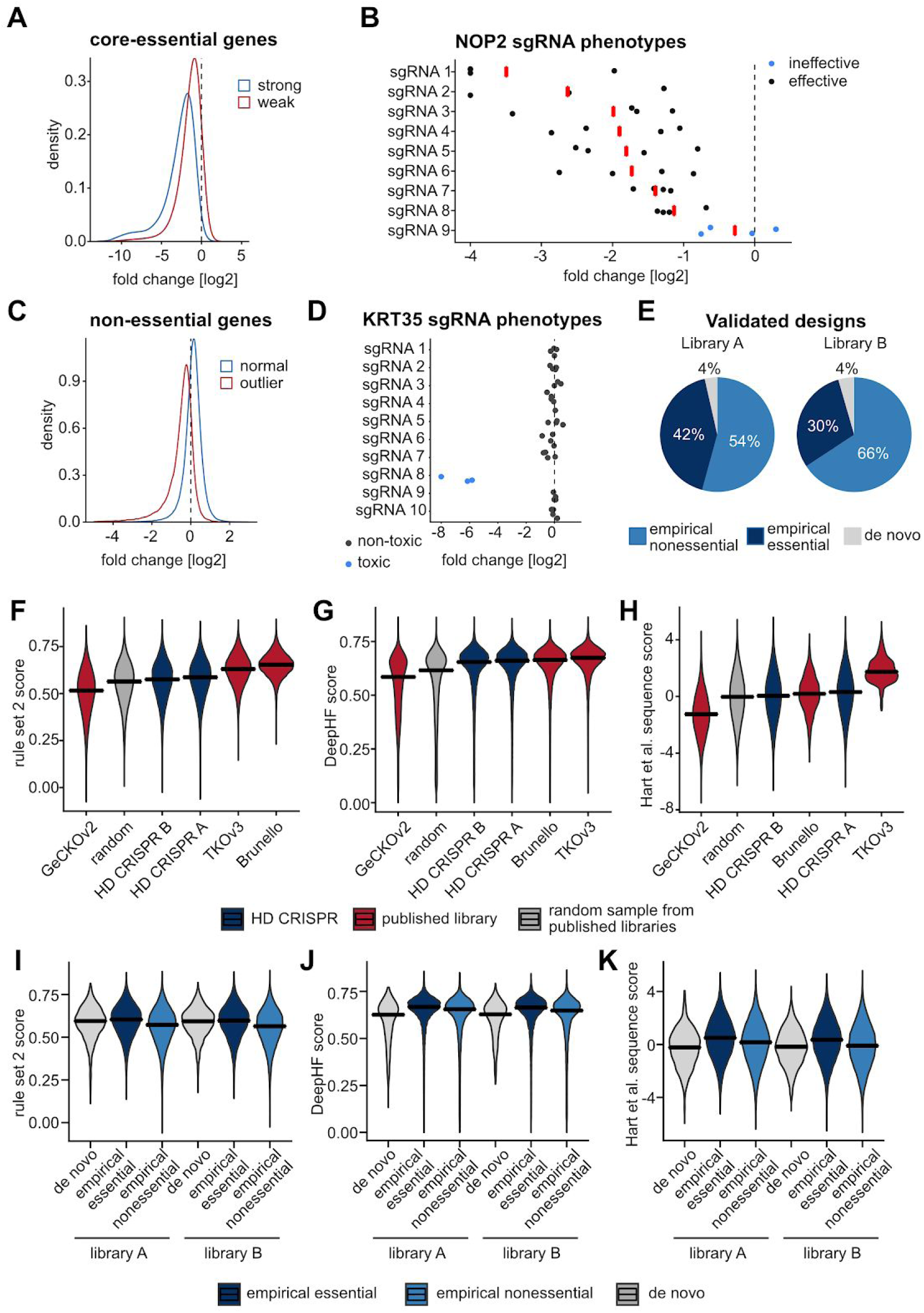
Empirically selected sgRNAs in the HD CRISPR Library. (A) Distribution of log2 fold changes for sgRNAs targeting core essential reference genes. The blue curve represents sequences with high on-target activity. The red curve represents sgRNAs that led to unexpectedly weak phenotypes. (B) Phenotypes of sgRNAs targeting the core essential gene NOP2 for 4 screens performed with the library described in Wang *et al.*, 2015 (Wang et al. 2015). Compared to other NOP2-targeting sgRNAs, sgRNA 9 showed unexpectedly weak depletion in these experiments and was thus labeled ‘ineffective’. (C) Distribution of log2 fold changes for sgRNAs targeting nonessential reference genes. The blue curve represents sequences with low off-target activity. The red curve represents sgRNAs that led to unexpectedly strong phenotypes. (D) Phenotypes of sgRNAs targeting the nonessential gene KRT35 in 4 screens performed with the library described in Wang *et al.*, 2015 (Wang et al. 2015). Unlike other KRT35-targeting sgRNAs, sgRNA 8 consistently displayed toxic phenotypes in these experiments and was therefore marked as ‘toxic’. (E) Percentage of sgRNA sequences in sub-libraries A (left) and B (right) that could be selected based on empirical evidence from published screening experiments. Empirical essential sgRNAs were selected based on inducing a viability phenotype, empirical nonessential sgRNAs based on the absence of a toxic phenotype. (F-H) Calculated sequence scores applying either the rule set 2 (Doench et al. 2016), the DeepHF (Wang et al. 2019) or the Hart et al. (Hart et al. 2017) algorithms. Score performance of the HD CRISPR sub-libraries A and B was benchmarked against the libraries whose design is based on respective scores (Brunello for rule set 2, TKOv3 for Hart et al.) if available as well as the GeCKOv2 library and a random sample of sgRNAs from published libraries. The DeepHF score was used as an independent measure none of the investigated libraries was designed on. (I-K) Comparison of sgRNA scores for empirically and de novo designed sgRNAs within the HD CRISPR sub-libraries.

Next, we identified and excluded sgRNAs with gene-independent toxic phenotypes (Doench et al. 2016). GenomeCRISPR provides an ‘sgRNA effect’ score that indicates how strong an observed sgRNA phenotype was in comparison to all other sgRNAs in the same screen (Rauscher et al. 2017a). We grouped sgRNAs by their target genes and scaled and centered their effect-scores to identify sgRNAs whose phenotypes strongly deviated from the phenotypes of other sgRNAs targeting the same gene (|sgRNA effect z-score| > 1.25). We labelled these sgRNA sequences as potentially off-targeting and excluded them from the library design (Figure 2 C-D). When less than 8 sgRNAs meeting these criteria were available from published libraries to target a gene, we used the CRISPR library designer (CLD; (Heigwer et al. 2016)) to design new sgRNAs (Supplementary Figure 1 C).

We then selected sgRNAs for a new library, named the Heidelberg (HD) CRISPR Library (Figure 1; Supplementary Table 1). This library consists of two independent, mutually exclusive sub-libraries A and B that each contain 4 sgRNAs per gene, targeting 18,913 and 18,334 protein coding genes, respectively (Supplementary Files 1 and 2). For selection, we prioritized sgRNAs that showed high on-target activity (as determined above) in a large number of screens.

If no information about sgRNA on-target activity was available (e.g. for nonessential genes), we picked sgRNAs that target constitutive exons with little predicted off-target effects (see Methods) close to the transcription start site (Supplementary Figures 1 D-E). Genes and their respective sgRNAs were categorized as essential if respective sgRNAs got depleted in at least 5% of the screens analyzed.

42% of the sgRNA for libraries A and 30% of the sgRNAs for library B met these criteria. An additional 54% of sgRNAs for library A and 66% of sgRNAs for library B could be empirically selected based on their uniform phenotype for presumably mainly nonessential genes. For only 4% of sgRNAs, *de novo* design was necessary (Figure 2E). Sub-library A contains sgRNAs that ranked best according to our design criteria to enable high-quality screens at a low library coverage. Sub-library B contains second-tier sgRNAs and can be used to supplement sub-library A when a higher sgRNA coverage is desired.

We further aimed to benchmark sgRNAs selected for the HD CRISPR Library based on commonly used sgRNA design scores. For comparison, we used sgRNAs from the GeCKOv2, TKOv3 and Brunello libraries and also generated a sample of 70,000 randomly selected sgRNAs from published libraries (Wang et al. 2014; Wang et al. 2015; Wang et al. 2017; Tzelepis et al. 2016; Sanjana et al. 2014; Doench et al. 2016; Doench et al. 2014; Hart et al. 2015; Hart et al. 2017; Munoz et al. 2016). For scoring, we applied the rule set 2 design rules, based on which the Brunello library was designed (Doench et al. 2016), as well as the sequence score developed for the design of the TKOv3 library (Hart et al. 2017). As an independent metric, we also evaluated sgRNAs using the more recently published DeepHF score (Wang et al. 2019), an sgRNA activity prediction score based on deep learning algorithms, which has not set the basis for the design of either of the evaluated libraries. For each of the selected scores, the two HD CRISPR sub-libraries outperformed the GeCKOv2 library and the randomly picked sample of published sgRNAs (Figure 2 F-H). The performance for the rule set 2 and Hart et al. sequence score were slightly lower for the HD CRISPR library (median 0.59 and 0.58 for sub-libraries A and B, respectively) in comparison to the Brunello and TKOv3 libraries (median 0.65 for Brunello and 0.63 for TKOv3). Since the HD CRISPR Library has solely been designed by relying on sgRNA associated phenotypes but not by considering any of these design rules, this is not surprising. However, using the independent deep learning DeepHF scoring system, all three empirically designed libraries (Brunello, HD CRISPR and TKOv3) performed similar and outperformed GeCKOv2 and the randomly picked sample (median scores: HD-CRISPR A = 0.66; HD-CRISPR B = 0.66; Brunello = 0.66; TKOv3 = 0.67; GeCKOv2 = 0.59; Random = 0.62) (Figure 2 G). Interestingly, sgRNAs empirically selected based on the induction of a viability phenotype scored better than sgRNAs selected for nonessential genes as well as *de novo* designed sgRNAs, for which only common design rules were applied (Figure 2 I-K).

### Cas9-editing efficiency can be improved by pre-selection of highly-editing single cell clones

Next, we further addressed how Cas9 activity could be increased to improve the sensitivity of CRISPR screens. We asked whether knockout efficiency could be enhanced in large screening settings by pre-selecting single cell clones with high Cas9 editing efficiency. Bulk populations of transduced cells are often heterogeneous in their transgene expression, largely due to differences in lentiviral integration, epigenetic silencing of exogenous DNA or the selection process during transfection or transduction (Kaufman et al. 2008). This heterogeneity can impact downstream processes such as Cas9-editing efficiency. In order to test for differences between bulk and clonal cells, we sorted single cells from the HAP1 Cas9 bulk population and measured Cas9 editing efficiency for several clones (Figure 3 A). From these, we selected two single cell clones, henceforth referred to as SCC11 and SCC12, with high Cas9 editing activity.

**Figure 3:**
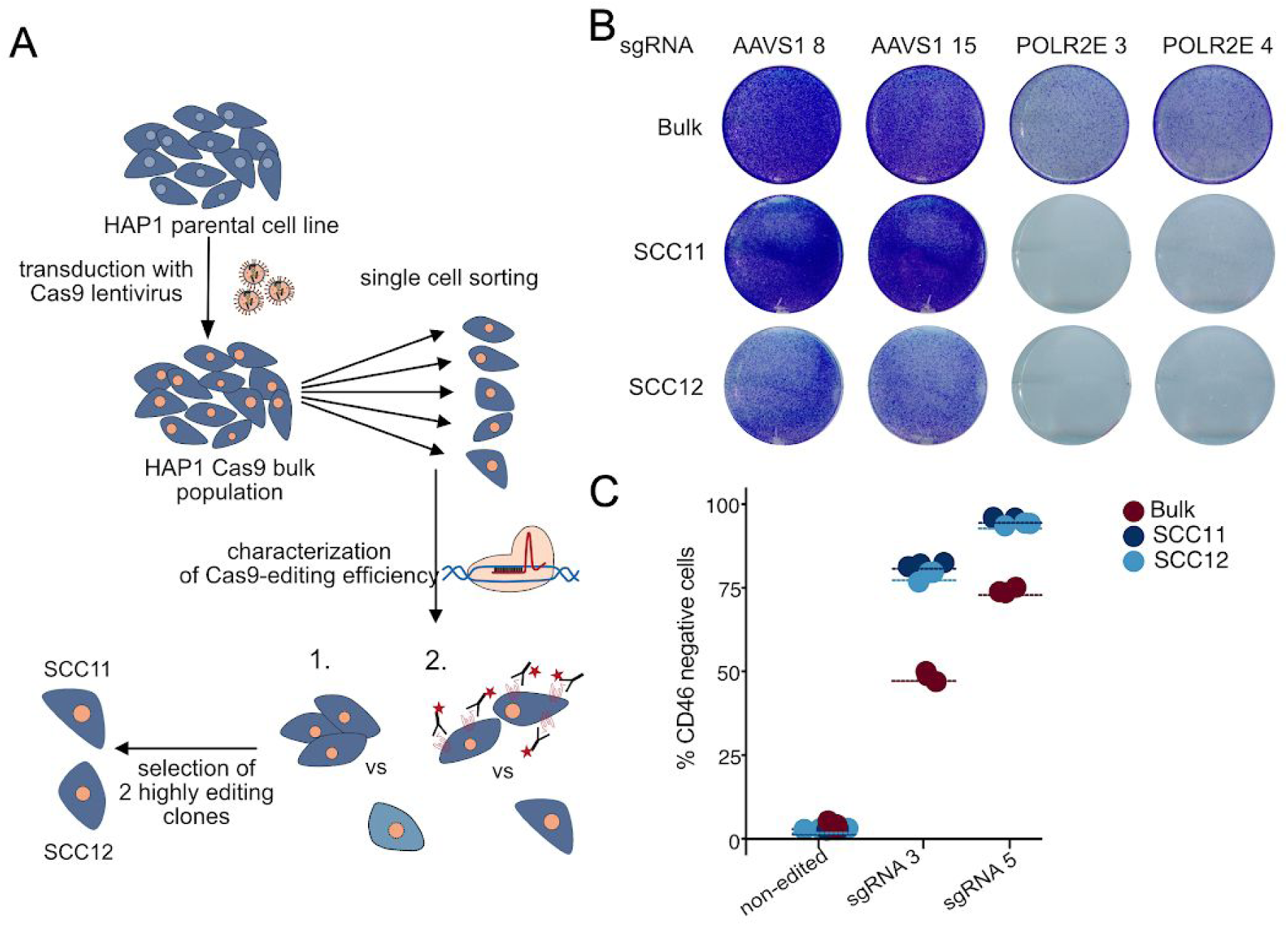
Selected Cas9-expressing single cell clones show stronger editing efficiency compared to a Cas9 bulk population. (A) Workflow for the selection of Cas9 single cell clones (SCCs). SCCs were sorted from the HAP1 Cas9 bulk population and further characterized. Cas9-editing was assessed by cell surface marker knockout followed by FACS staining and cell viability upon knockout of a core essential gene. Two highly editing single cell clones (SCC11 and SCC12) were selected for further experiments. (B) HAP1 Cas9 bulk, Cas9 SCC11 and Cas9 SCC12 cells were transfected with the HD CRISPR vector encoding an sgRNA targeting either the safe harbour locus *AAVS1* as a control or the core essential gene RNA Polymerase 2 subunit E (*POLR2E*). Editing efficiency based on cell viability of sgPOLR2E transfected cells in comparison to sgAAVS1 control cells was addressed by crystal violet staining. The number of surviving cells was strongly reduced in cells transfected with an sgRNA directed against *POLR2E.* (C) Editing efficiency was furthermore assessed upon transduction of HAP1 Cas9 bulk, Cas9 SCC11 and Cas9 SCC12 cells with the HD CRISPR vector expressing sgRNAs targeting the surface marker *CD46*, followed by FACS staining of residual CD46 protein to address knockout efficiency. Antibody staining of the non-edited cell lines was used as a control.

For our knockout experiments, we designed an sgRNA expression vector with an improved sgRNA scaffold (Dang et al. 2015; Tzelepis et al. 2016). The vector furthermore features a stuffer sequence encoding GFP and a lentiviral backbone similar to the previously published pLCKO vector (Hart et al. 2015). Cloning of sgRNA sequences results in removal of the GFP stuffer (Supplementary Figure 2 A), which allows to estimate the extent of remaining non-digested vector backbone in library preparations and thus cloning efficiency (Supplementary Figure 2 B-C), while upon transduction the stuffer does not interfere with assays where GFP is used as a readout (Supplementary Figure 2 B). The functionality of the HD CRISPR vector was confirmed by knockout of the surface marker gene *CD81* in HAP1 Cas9 bulk cells (Supplementary Figure 2 D, Supplementary Table 2).

Using this vector to compare knockout efficiency in Cas9 bulk and single cell clones, knockout of the core essential *POLR2E* gene led to much stronger depletion of viable cells in SCC11 and SCC12 compared to the Cas9 bulk population (Figure 3 B). Similarly, when addressing the amount of remaining surface protein upon knockout of *CD46* using two individual guides, depletion was 20% to 30% stronger in single cell clones compared to the bulk population (Figure 3 C; Supplementary Table 2). We considered it possible that editing efficiency might be affected by the ploidy of the different cell lines. While HAP1 cells are a haploid cell line (Carette et al. 2011), their haploid state tends to be unstable. In a mixed population of haploid and diploid HAP1 cells, diploid cells have been shown to enrich over time due to a proliferative advantage, while the haploid state can be prolonged in clonal populations starting from a single haploid cell (Olbrich et al. 2017). In line with this, we identified a larger proportion of clearly diploid cells in the HAP1 Cas9 bulk population compared to the two single cell clones (∼11.5% diploid in HAP1 Cas9 bulk vs. ∼1.5% in HAP1 Cas9 SCC11 and ∼6.3% in HAP1 Cas9 SCC12) (Supplementary Figure 3 A). To rule out that ploidy has a major impact on editing efficiency in the HAP1 cell line, we sorted enriched haploid (68.2% for SCC11 and 66.8% for SCC12) and diploid populations (43.2% and 34.2% for SCC11 and SCC12, respectively) from the HAP1 Cas9 SCC11 and SCC12 cell lines. Subsequently, we directly compared editing efficiency in enriched haploid and diploid populations originating from the same single cell clone. While we indeed observed a slightly lower editing efficiency in diploid cells compared to haploid controls, this difference was smaller than 6% in all cases (Supplementary Figure 3 B-C). This argues for other factors than ploidy to be the main drivers of differences in editing efficiency.

### The HD CRISPR Library identifies core and nonessential genes at high precision

In order to address screening performance of the HD CRISPR Library, the sub-libraries A and B were cloned independently into the corresponding HD CRISPR vector. Quality controls of the resulting plasmid preparations revealed a narrow sgRNA distribution with negligible background of non-digested vector backbone (Supplementary Figure 4). Both libraries were screened in the HAP1 Cas9 bulk population and the two Cas9 single cell clones in two independent replicates for 14 days to further address the impact of Cas9 editing efficiency and the extent of clonality effects on hit calling (Figure 4 A; Supplementary Figure 5). Since our reference set of published CRISPR screens used for library design did not include any screen conducted in HAP1 cells, we considered this cell line to be a suitable model to address library performance.

**Figure 4:**
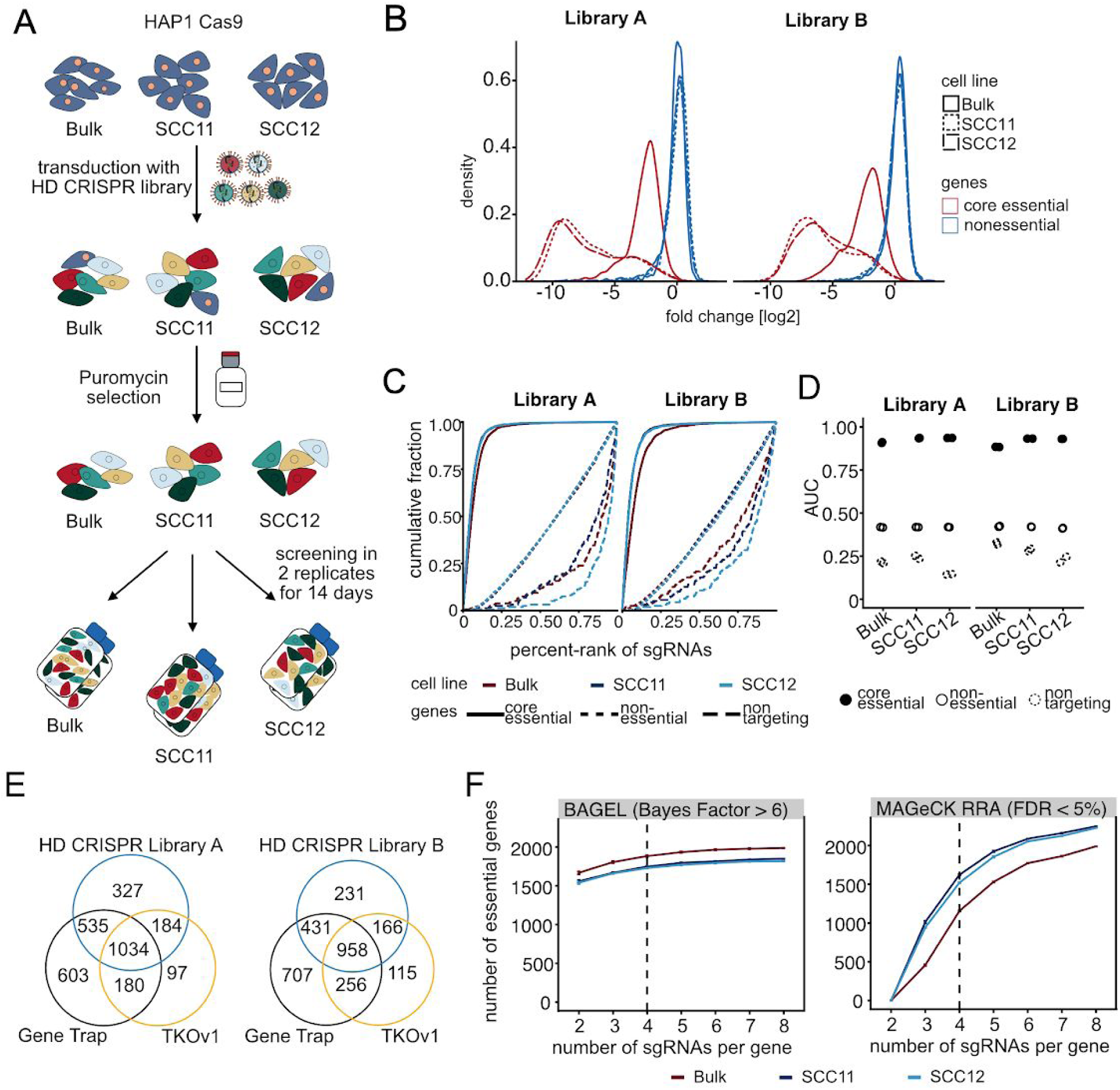
The HD CRISPR Library efficiently identifies core, non and context-dependent essential genes. (A) Workflow of a pilot screen conducted with the HD CRISPR Library in HAP1 cells. The screen was performed in parallel in the Cas9-expressing bulk population and two highly-editing single cell clones for both libraries independently. Successfully transduced cells were selected with puromycin for 48 h and then split into two independent replicates. The screen was performed for a duration of 14 days. (B) Core essential genes were strongly depleted over the course of screening with either of the two libraries, HD CRISPR Library A and B, in contrast to nonessential genes. Stronger depletion was observed in the two single cell clones with high Cas9 editing efficiency. (C) Area under the curve (AUC) of the precision recall curve for viability screens conducted with each HD CRISPR sub-library in the HAP1 Cas9 bulk, Cas9 SCC11 and Cas9 SCC12 cell line for core essential, nonessential and non targeting gene sets. (D) Comparison of AUCs for individual replicates of screens conducted in each HAP1 Cas9 cell line using each of the HD CRISPR sub-library. (E) Comparison of HAP1 essential genes as identified with the HD CRISPR Library, a gene trap screen by Blomen et al. (2015) and a CRISPR screen using the TKOv1 library by Hart et al. (2017). (F) Number of essential genes detected with increasing number of sgRNAs per gene using BAGEL (left; BF > 6) or MAGeCK RRA (right; FDR < 5%). sgRNAs were subsampled from the combined HD CRISPR Library. Each data point represents the average of 5 samples. Error bars are ± 1 s.e.m.

Comparing fold changes of sgRNAs targeting genes comprising a core essential and a nonessential reference gene set (Hart et al. 2014, Hart et al. 2017) confirmed strong loss of sgRNAs targeting essential genes over the course of screening for both libraries and all cell lines, while the representation of nonessential genes remained nearly unchanged to the plasmid library, validating our sgRNA selection strategy (Figure 4 B; Supplementary Table 3). As expected, depletion of sgRNAs targeting essential genes was stronger in selected single cell clones compared to the bulk population and screening in single cell clones furthermore strongly improved replicate correlation in comparison to screening in HAP1 Cas9 bulk cells (Supplementary Figure 5). This confirms our assumption that a better resolution can be achieved upon screening in selected single cell clones pre-selected for strong Cas9 editing efficiency.

To further address library performance, we computed the area under the precision-recall curve (AUC) using core essential, nonessential and non targeting reference gene sets (Hart et al. 2014; Hart et al. 2017) (Figure 4 C-D). Effective screens are supposed to have an AUC value ≥ 0.5 for sgRNAs targeting essential genes and an AUC value ≤ 0.5 for sgRNAs targeting nonessential genes or designed as non targeting controls (Sanson et al. 2018) indicating that respective sgRNAs either get preferentially depleted (AUC ≥ 0.5) or remain (AUC ≤ 0.5) over the course of screening. We obtained AUC values close to 1 for both independent replicates of all screens when analyzing sgRNAs targeting essential genes, and AUC values below 0.5 for sgRNAs designed for nonessential genes and as non targeting controls (Figure 4 D), indicating that our selection strategy for core and nonessential genes was successful.

To benchmark our results with other screens conducted in HAP1 cells, we compared genes identified to be essential in the HAP1 Cas9 bulk population using either the HD CRISPR sub-libraries A or B with published data from gene essentiality screen conducted in HAP1 cells using either the TKOv1 library (Hart et al. 2015) or a gene trap screen (Blomen et al. 2015). The intersection of hits from all three screens was greater than 900 genes for both HD CRISPR sub-libraries (Figure 4 E) and hits identified in our screens largely corresponded to those identified in previous screens (Supplementary Figure 8 A and B), suggesting that hits from previous screens - including HAP1 context-dependent essential genes - can be re-discovered using the HD CRISPR Library.

We furthermore compared hit calling of essential genes using different analysis methods, BAGEL and MAGeCK RRA. Using BAGEL, differences in the number of essential genes identified were marginal for both, increasing numbers of sgRNAs per gene (2-8) and screening in either the Cas9 bulk population or one of the two single cell clones (Figure 4 F). This is likely due to the fact that BAGEL uses prior knowledge for gene essentiality hit calling. In contrast, for analysis softwares, which do not require prior information such as MAGeCK RRA or gscreend (Li et al. 2014; Imkeller et al. 2020), an increasing number of sgRNAs per gene also allowed the identification of more hits and single cell clones with an enhanced editing efficiency were superior for hit calling in comparison to the bulk population (Supplementary Figure 6 A-C). Overall, comparing genes identified to be essential using either BAGEL or MAGECK RRE revealed strong agreement with few essential genes private to each software (Supplementary Figure 6 D).

In a final step, we asked how differences in editing efficiency, the number of sgRNAs per gene as well as potential clonality effects might affect the determination of gene essentiality. We were especially interested in differences in gene essentiality between the Cas9 bulk cell line and the two Cas9 single cell clones populations, since the generation of single cell clones from bulk populations forces cells to go through a genetic bottleneck, which might favor certain genetic alterations (Giuliano et al. 2019) and thus dependencies. Therefore, we used BAGEL (Hart and Moffat 2016) to analyze depleted genes in each HAP1 cell line using the combined HD CRISPR library (8 sgRNAs per gene) and the individual sub-libraries A and B. We applied a strict Bayes Factor cutoff of BF > 6 (Hart et al. 2017) to discriminate between essential and nonessential genes. This analysis revealed highly overlapping sets of essential genes for each cell population. Using the combined library we found 2096 essential genes in the Cas9 bulk population, 1938 essential genes in SCC11 and 1925 essential genes in SCC12. Out of these, 1755 were shared between the three lines (84%, 91% and 91% of total essential genes in each cell line). Only 58 (3% of total) essential genes were private to SCC11 and 46 (2.5% of total) essential genes were private to SCC12 (Supplementary Figure 7A). We observed similar overlap using only sub-library A or B (Supplementary Figure 7C and D). Accordingly, quantitative comparison revealed high Bayes Factor correlation between bulk Cas9 population and single cell clones (0.915 for SCC11 and 0.925 for SCC12; Supplementary Figure 7 B). Overall, we could not observe major differences in hit calling between selected Cas9 single cell clones and a bulk population when addressing general gene essentiality. We, however, think that caution is required when screening in more complex settings as e.g. when addressing drug resistance mechanisms.

### CRISPR screens at high sensitivity allow predictions about sgRNA cutting efficiency

CRISPR-induced DNA double strand breaks at target loci are known to induce a DNA damage response, which significantly affects cell proliferation in comparison to negative controls (Aguirre et al. 2016). As a result, non-targeting sgRNAs are likely to reflect wild type growth, while sgRNAs targeting nonessential genes and targeting controls lead to slightly impaired growth (Hart et al. 2017). Differences between targeting and non-targeting controls are usually subtle. However, small but distinct shifts in the fold change distributions of non-targeting and targeting control sgRNAs were visible in all our screens in the HAP1 cell line (Supplementary Figure 8A). This shift was especially pronounced in the screens conducted in the HAP1 Cas9 SCC12 cell line (Figure 5A, Supplementary Figure 8A). We reasoned that this effect could be explained by the increased sensitivity that can be achieved by using a single cell clone selected for high Cas9 activity. We hypothesized that it might be possible to exploit this increased resolution to assess the activity of sgRNAs targeting nonessential genes. Using a Gaussian mixture model (Benaglia et al. 2009) we divided all control sgRNAs in the SCC12 screen into three groups: (1) sgRNAs with strong viability phenotypes, (2) sgRNAs targeting nonessential loci (mild phenotype due to induction of DNA double strand breaks) and (3) inactive sgRNAs (Figure 5A; Supplementary Figure 8B-C). We then classified all sgRNAs in the HD CRISPR Library into one of these groups. To avoid bias we excluded reference core essential genes (Hart et al. 2017) from the analysis. sgRNAs that could not be assigned to any of the groups with a probability of at least 80% were labeled ‘undetermined’ (Figure 5A).

**Figure 5:**
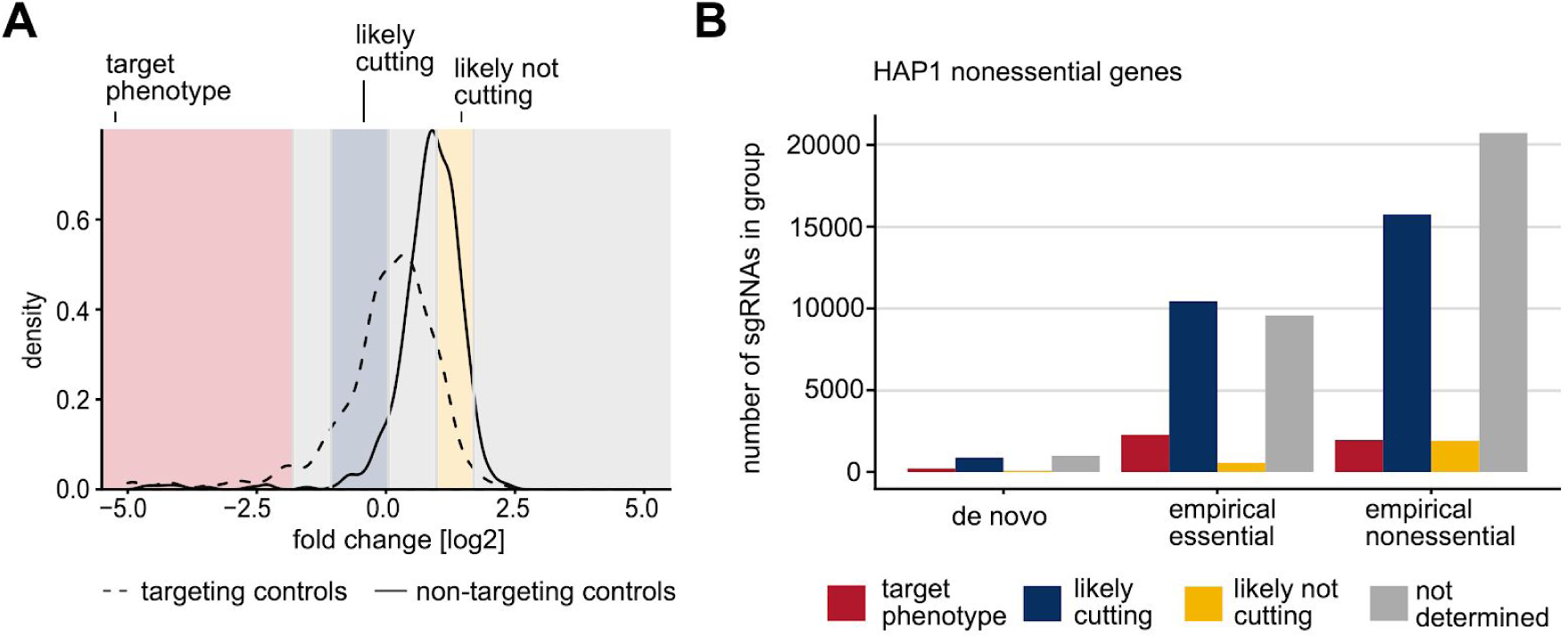
CRISPR screens conducted at a high dynamic range predict the cutting efficiency of sgRNAs based on mild viability phenotypes. (A) A mixture model was used to divide control sgRNAs of HD CRISPR Library A in the single cell clone SCC12 into three groups: (1) sgRNAs with a target-dependent viability phenotype (red), (2) sgRNAs with a small target-independent phenotype likely caused by a double strand break (blue), and (3) sgRNAs with no phenotype due to a lack of DNA cutting (yellow). Log2 fold change distributions of targeting and non-targeting control sgRNAs are indicated as dashed and solid curves, respectively. B) Number of sgRNAs targeting nonessential HAP1 genes associated with each phenotype group. Nonessential genes were determined using MAGeCK which requires no prior knowledge for analysis. sgRNAs are stratified based on their design: ‘empirical essential’ sgRNAs target context-specific essential genes and were selected for the HD CRISPR Library based on their previous on-target phenotypes. ‘Empirical nonessential’ sgRNAs are part of previously published libraries and target broadly nonessential genes. They were selected based on their lack of outlier phenotypes. De novo sgRNAs were designed using the software cld (Heigwer et al. 2016).

**Figure 6:**
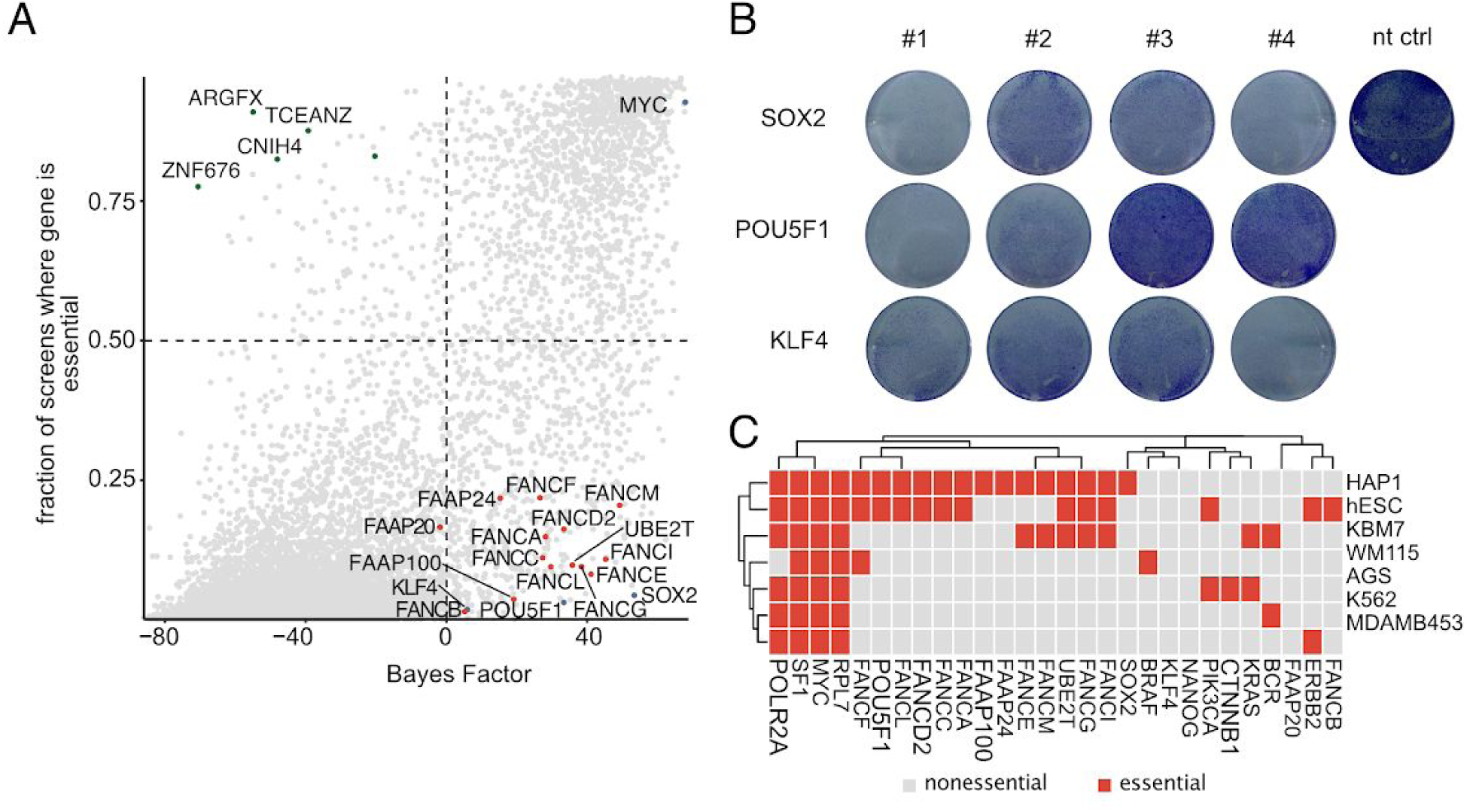
HAP1 cells are highly dependent on Yamanaka factors and the Fanconi anemia pathway. (A) HAP1 context-dependent essential genes are enriched for genes comprising the Fanconi anemia pathway as well as factors known to induce pluripotency. The y-axis provides an estimate of the percentage of essentiality in other cell lines previously screened. Fanconi anemia pathway genes are highlighted in red, Yamanaka factors are shown in blue. HAP1 specific nonessential genes are highlighted in green. (B) Crystal violet staining of HAP1 cells treated with various siRNAs targeting *SOX2, POU5F1* and *KLF4* and a non-targeting control. (C) Essentiality of known cancer drivers, Fanconi anemia pathway genes and pluripotency factors across cell lines representing different cancer cell types in comparison to HAP1 cells and human embryonic stem cells.

As expected, most sgRNAs targeting essential genes (determined using MAGeCK analysis (Li et al. 2014) of the combined HD CRISPR Library) were classified as group 1. This effect was more pronounced for sgRNAs selected based on previous phenotypes compared to other sgRNAs (82% and 60% of sgRNAs targeting essential genes classified as group 1, respectively; Supplementary Figure 8D). In addition, most sgRNAs targeting nonessential genes were classified as group 2 (likely cutting) whereas only a small fraction of sgRNAs were classified as group 3 (likely not cutting). Again, sgRNAs selected based on their phenotypes in previous screens appeared favorable compared to other sgRNAs with a larger fraction classified as likely cutting (46% compared to 39%) and a smaller fraction classified as likely not cutting (2% compared to 5%; Figure 5B; Supplementary Figure 8E). These observations suggest that sgRNA phenotypes in past screens are in fact predictive of whether these sgRNAs will be effective in future screens and thus motivate empirical design as a viable strategy for sgRNA selection - not only for core essential genes but also for context-specific essential genes.

### Identification of HAP1 context-dependent essential genes

Selecting sgRNAs with consistent knockout phenotypes across several independent screens is most conclusive for broadly essential genes. Therefore, we examined whether the HD CRISPR Library could also identify context-dependent essential genes and as such genes that are specifically essential in HAP1 cells. We compared the list of hits identified in our screens to essential genes reported in other screens and cell lines in the GenomeCRISPR database. We identified a set of genes (including *ARGFX, CNIH4, TCEANC* and *ZNF676*), which appeared to be essential in more than 75% of published screens we used for analysis, but not in our screens in HAP1 cells using the HD CRISPR library (Figure 5 A). These might reflect genes essential in most cell lines, but not HAP1, or might reflect genes that have been frequently wrongly annotated as essential. When comparing Bayes Factors (BF) as a measurement for gene essentiality for these genes across different libraries, we observed that these genes were identified as essential in screens with the Avana sgRNA library (Doench et al. 2014), but not with other published genome-scale libraries (Supplementary Figure 9). This suggests that the unexpected and broadly observed essentiality of these genes might reflect artefacts associated with the respective sgRNAs present in the Avana library. Such off-targets are likely found in each CRISPR library and only become apparent when many different cell lines are screened with the same library as is the case for the Avana library. Importantly, this implies that our selection strategy, although based on choosing sgRNAs with strong phenotypes in many screens, does not enrich for sgRNAs with strong off target effects.

While the near-haploid karyotype of HAP1 cells renders them particularly amenable for genetic perturbation studies (Blomen et al. 2015), relatively little is known about their identity and cell line-specific dependencies. Thus, we addressed HAP1 context-dependent essential genes as identified in our screen in more detail. In general, roughly 2000 genes are considered to be essential in cultured human cells (Rancati et al. 2018), which is in accordance with our results (Figure 4 F). With ∼700 genes comprising the core essential gene set (Hart et al. 2017), ∼1300 of context-dependent essential genes can be considered to be identified. HAP1 cells originate from experiments to induce pluripotency in the leukemia KBM7 cell line by transduction with *KLF4, POU5F1* (Oct4), *SOX2* and *MYC* (Carette et al. 2011; Takahashi and Yamanaka 2006). *Interestingly, our screening results indicate that this treatment seemed to have rendered HAP1 cells strongly dependent on POU5F1* and *SOX2* and to a lesser extent also on *KLF4.* In addition, we identified genes comprising the core complex of the Fanconi anemia pathway to be essential (BF > 6) in HAP1 cells, while often dispensable in other cell lines (Figure 5 A; Supplementary Figure 10 C).In contrast, *KLF4, POU5F1 and SOX2* were only identified to be essential in less than 5% of other cell lines screened so far (Figure 5 A). These genes were not identified as essential in former gene trap screens in HAP1 cells (Supplementary Figure 10 D), which might be explained by the fact that HAP1 cells likely carry several copies of *KLF4, POU5F1* and *SOX2* due to their initial transduction. Hence, we further wanted to exclude that the observed essentiality was due to multiple integrations of the lentiviral expression vector and thus an increased DNA damage response upon several CRISPR-induced cuts (Aguirre et al. 2016; Munoz et al. 2016). Therefore, we validated the dependency of HAP1 cells on *KLF4, POU5F1* and *SOX2* using siRNAs as an orthologous approach (Figure 5 B, Supplementary Table 4).

The dependency on pluripotency factors prompted us to compare gene essentiality of HAP1 cells with previously published CRISPR data for gene essentiality of the original KBM7 cell line, as well as of human embryonic stem cells (hESC) (Wang et al. 2015; Yilmaz et al. 2018). The comparative analysis revealed that loss of *POU5F1* and some of the components of the Fanconi anemia core complex also resulted in reduced cell viability in hESCs. In contrast, KBM7 cells displayed a dependency on several genes associated with the Fanconi anemia pathway, but not for either of the Yamanaka factors except for the core essential gene *MYC* (Figure 5 C). Interestingly, viability of HAP1 cells did not seem to be affected by the knockout of known tumor type-specific oncogenes such as *BCR, PIK3CA, KRAS, CTNNB1* or *BRAF* (Figure 5 C). Especially the lack of BCR dependency is surprising, since both HAP1 cells and their parental cell line KBM7, harbor a BCR-ABL fusion gene, which, however, is only required for proliferation in KBM7 cells. Based on their shared dependencies with hESCs while missing an addiction to known tumor type-specific oncogenes, we suggest that the initial transduction with pluripotency factors has rendered HAP1 cells to adopt stem cell-like characteristics.

## DISCUSSION

Over the last few years, pooled genome-scale CRISPR-Cas9 screens have quickly become an established and indispensable tool to functionally interrogate the human genome. Still, much remains to be learnt about optimal experimental conditions and sgRNA design to optimize especially the dynamic range and phenotype detection in CRISPR screens, while minimizing expenses and workload.

To date, large-scale CRISPR-Cas9 screens have been conducted in hundreds of human cell lines (Wang et al. 2015, 2017; Hart et al. 2015; Aguirre et al. 2016; Meyers et al. 2017; Behan et al. 2019). These data now enable us to draw conclusions about the efficacy of hundreds of thousands of sgRNAs in diverse experimental contexts. Here, we introduced a strategy to leverage data from published CRISPR screens for sgRNA design. Our goal was to prioritize sgRNAs with consistently high on-target activity, while simultaneously avoiding sgRNAs with off-target effects. We were able to select high-quality sgRNAs and to assemble them into the Heidelberg (HD) CRISPR library. More than 40% of sgRNAs for both sub-libraries were selected based on empirical essentiality phenotypic evidence, while we could select non-toxic guides for the remaining 50 to 60% of protein-coding genes, generating libraries with overall 96% of empirically designed guides. This number is limited primarily by the fact that currently available CRISPR screening data are mostly derived from viability screens for gene essentiality. Therefore, we expect this number to grow quickly once screens for additional phenotypes become available.

We could show that guides in this library were enriched for high sequence scores according to different design rules although these were not initially taken into account for library design. To evaluate the performance of this library we conducted a genome-wide screen for gene essentiality in HAP1 cells and confirmed that our library was able to distinguish between core and nonessential reference genes with high precision and accuracy. Since it is desirable to minimize library size for CRISPR screens in order to reduce expenses and experimental efforts, we assessed how the number of sgRNAs per gene would affect the detection of essential genes. Therefore, we compared screens with the combined HD CRISPR Library (8 sgRNAs per gene) and screens in individual sub-libraries A and B (4 sgRNAs per gene). Screening with only one of the two sub-libraries reduces experimental efforts by ∼ 3.5*10^7^ cells per replicate and split when screening with a 500-fold coverage.

To further analyze how these results would be affected by editing efficiency and clonality effects, we performed screens in a HAP1 Cas9 bulk population as well as two single cell clones. Overall, essential genes were highly consistent between the bulk population and the single cell clones, while phenotypes were persistently stronger in the single cell clones (Figure 3 C, Figure 4 B). This could not be explained by differences in ploidy of Cas9 single cell clones and the bulk population, since enriched diploid populations of the two HAP1 Cas9 single cell clones still showed similar editing efficiency to enriched haploid populations and remained superior in comparison to Cas9 bulk populations (Supplementary Figure 3 B-C). This indicates that interpopulation heterogeneity is a stronger determinant of editing efficiency than ploidy. While it is advisable to screen and compare more single cell clones also in different cellular contexts to draw definite conclusions, our results suggest that single cell clones with high Cas9 activity are attractive models to screen for gene essentiality at small library sizes. While we did not observe strong clonality effects between single cell clones for detecting essential genes, it is likely that other phenotypes might be affected more severely by clonal differences. Therefore, caution is especially required when using single cell clones in screens for phenotypes such as pathway activity or cell morphology. A middle-way could be to pool several independent single cell clones with similar proliferation rates into a pseudo bulk population that can be used for screens. By this means, clonality effects may be minimized while still maintaining high editing efficiency.

Haploid cell lines such as HAP1 cells are an attractive model for genetic perturbation studies since the presence of a single allele implies enhanced knockout efficiency (Blomen et al. 2015). Although we could show that knockout efficiency is only slightly decreased in diploid compared to haploid HAP1 cells, the effect is considered to be stronger in polyploid cancer cell lines where a full knockout requires out-of-frame editing of all alleles of a gene (Van Campenhout et al. 2019). HAP1 cells with specific gene knockouts have therefore been applied in many studies addressing various research questions (Davis et al. 2015; Schick et al. 2019; Lenk et al. 2019; Smits et al. 2019; Gerhards et al. 2018; Baggen et al. 2019). The exact cellular context of this cell line is, however, not fully understood, especially since HAP1 cells do not share major characteristics with their parental chronic myeloid leukemia (CML) cell line KBM7. In contrast to KBM7 cells, HAP1 cells grow adherent and are not dependent on *BCR* (Figure 5 C), while Bcr-Abl inhibitors represent the first-line therapy in CML (Rossari et al. 2018). When specifically addressing HAP1 context-dependent gene essentiality, we identified known and so far unreported HAP1 context-specific essential genes. In particular, we describe a strong HAP1-specific dependency on the Yamanaka factors *POU5F1, SOX2* and to a lesser extent also *KLF4* (Takahashi and Yamanaka 2006). These genes have been transduced into KBM7 cells during the generation of the HAP1 cell line (Blomen et al. 2015). Further, HAP1-specific essential genes were enriched for components of various complexes of the Fanconi anemia pathway. The Fanconi anemia pathway is mainly known to be responsible for the repair of stalled replication forks that occur as a consequence of interstrand crosslinks (Ceccaldi et al. 2016). Recently it has been reported that *FANCD2* localizes to sites of Cas9-induced DNA double strand breaks where it supports CRISPR-mediated homology directed repair and in particular single-stranded template repair (Richardson et al. 2018). Interestingly a dependency on certain Fanconi anemia regulators can also be observed in two other haploid cell lines. These include the near-haploid HAP1 parental cell line KBM7 (Wang et al. 2015) and a haploid human embryonic stem cell line (Yilmaz et al. 2018). Since regulators of the Fanconi anemia pathway are rarely essential in other cell lines, we speculate that the occurrence of interstrand crosslinks might be more detrimental in cells featuring only one copy of any given gene.

In conclusion, we show that the available data on sgRNA phenotypes in large-scale CRISPR essentiality screens can be used to inform the empirical design of sgRNA libraries. We provide the HD CRISPR Library, a new library for genome-wide CRISPR-Cas9 screens with high on-target and low off-target activity. We further show evidence that single cell clones are powerful models to conduct gene essentiality screens at a low library coverage of sgRNAs per gene and can improve screening quality over screening in Cas9 bulk populations. These findings might guide experiment design for the next generation of CRISPR-Cas9 screens in mammalian cell culture.

## CONCLUSIONS

We conclude that conducting CRISPR/Cas9 viability screens using next generation empirically designed sgRNA libraries and strongly-editing Cas9 single cell clones improves the resolution of CRISPR screens and to differentiate also subtle viability phenotypes, which in turn allows hit detection with a smaller number of sgRNAs per gene. We show that empirical design of sgRNA libraries based on phenotypic evidence from previous screens is a suitable predictor for sgRNA cutting efficiency and allows selection of highly active guides, which we assembled in a genome-scale CRISPR library termed “HD CRISPR Library”.

## MATERIALS AND METHODS

### Heidelberg CRISPR library design

Raw sgRNA count data were downloaded from the GenomeCRISPR database (November 5th, 2017; (Rauscher et al. 2017a). Only negative selection screens performed using humanized S. pyogenes Cas9 were retained for downstream analysis. At the time of sgRNA design, no published screens were available with the TKOv3 (Hart et al. 2017) and Brunello (Doench et al. 2016) libraries. Therefore, these sgRNA sequences were added to the GenomeCRISPR-derived sgRNA pool. Next, all targeted transcripts and protein coding exons were annotated for each sgRNA. For this purpose, genomic information was derived from the ENSEMBL database (GRCh38; (Cunningham et al. 2019) using biomaRt (Drost and Paszkowski 2017). In order to determine putative off-target effects for each sgRNA, sgRNA sequences were mapped to the genome using bowtie2 (Langmead and Salzberg 2012). Specifically, local alignment was performed using bowtie2’s ‘very-sensitive-local’ setting allowing for up to 3 mismatches. This strategy for computational off-target prediction was motivated by previous studies (Heigwer et al. 2014, 2016).

To additionally avoid sgRNAs with off-target effects, sgRNAs were grouped by their target genes and their GenomeCRISPR effect scores (Rauscher et al. 2017a) were z-normalized. This was done to identify sgRNAs whose phenotypes strongly deviate from the phenotypes of other sgRNAs targeting the same gene. Increasing depletion of sgRNAs targeting nonessential genes and lack of depletion of sgRNAs targeting core essential genes was detected at effect scores smaller than −1.25 and larger than 1.25, respectively (Supplementary Figure 1C). Therefore, sgRNAs with an absolute effect score of greater than 1.25 were flagged.

Next, sgRNA sequences containing consecutive stretches of the same nucleotide (4 or more A’s/T’s, 5 or more G’s/C’s) were flagged since these sequences have been shown to create problems during polymerase transcription and PCR amplification. In addition, sequences containing BbsI restriction sites (GAAGAC and reverse complement) and sequences with a strong GC bias (GC greater than 75% or less than 20%) were excluded. To determine sgRNA performance all negative selection (for viability) screens in GenomeCRISPR (in total 488) were analyzed using BAGEL v0.91 (Hart and Moffat 2016) with the CEGv2 and NEGv1 core and nonessential reference gene sets (Hart et al. 2014, 2017). To evaluate the quality of each screen, precision-recall curves (PR-curves) were generated for each experiment using the ROCR R package (Sing et al. 2005). These PR-curves evaluate how well reference core and nonessential genes could be separated based on the sgRNA depletion phenotypes in the screen. Screens for which the area under the PR-curve was less than 0.9 were excluded from further analysis. For all other screens, genes were categorized as essential and nonessential at 5% false discovery rate (FDR). An sgRNA was then determined as active in a screen if (a) it was determined to target an essential gene (in the screened cell line) and (b) its depletion phenotype (quantified as sgRNA count fold change compared to a T_0_/plasmid sample) was among the 20% strongest sgRNA phenotypes in the screen. In a genome-wide CRISPR screen for cell proliferation, approximately 12% of all protein coding genes are expected to be essential (Meyers et al. 2017; Dempster et al. 2019) and therefore 12% of all sgRNAs are expected to have an on-target phenotype. However, in order to not miss sgRNAs with potential subtle phenotypes, we selected a lenient threshold of 20% to determine active sgRNAs. sgRNAs for the HD CRISPR Library were finally selected from the remaining pool of sequences. sgRNAs that were determined as active in a large number of screens were prioritized. However, to avoid selecting sgRNAs based on spurious effects, sgRNA activity was only considered as selection criteria if the sgRNA was determined as active in at least 5% of the screens in which they were used. Otherwise, sgRNAs targeting exons present in many transcript isoforms, sgRNAs targeting exons close to the transcription start site and sgRNAs with a low number of predicted off-target effects were prioritized. In total a genome-wide library consisting of two mutually exclusive sub-libraries A and B was assembled. Each sub-library contains 4 sgRNAs per gene targeting 18,913 and 18,334 protein coding genes, respectively. In cases where less than 4 sgRNAs were available for a gene based on the filtered pool of sequences described above, missing sgRNAs were designed with the CRISPR library designer (Heigwer et al. 2016). Further 300 non-targeting controls derived from published libraries (Wang et al. 2014, 2015; Hart et al. 2015) and 135 targeting controls targeting intronic regions or the AAVS1 safe harbor locus (Heigwer et al. 2014) were added to each sub-libraries. Control sgRNAs are identical between the sub-libraries. In total this resulted in sub-libraries of sizes 74,987 (library A) and 71,048 (library B) that can be combined into a library of size 146,035 for increased detection power.

### Cloning of the HD CRISPR sgRNA vector

For generation of the HD CRISPR vector, an insert encoding the human U6 promoter, an eGFP stuffer, part of the improved sgRNA scaffold (Dang et al. 2015), the cPPT/CTS, the human PGK promoter, a puromycin N-acetyltransferase and the WPRE next to stuffer sequences was ordered as a GeneArt Synthetic Gene (Life Technologies) and PCR-amplified. The bacterial backbone of the pLCKO vector (kind gift from the Moffat lab, Addgene #73311) was linearized from the 3’LTR to the RRE by PCR and both PCR products were fused using In-Fusion Cloning (Takara Bio) and transformed into One Shot Stbl3 Chemically Competent E.coli (Life Technologies). A correct cloning product was identified by control restriction enzyme digestion using AarI (Thermo Fisher Scientific) and subsequent Sanger sequencing. The initial plasmid only contained half of the improved sgRNA scaffold due to different possible sgRNA cloning strategies. To clone the complete improved sgRNA scaffold inside, the plasmid was digested using AarI (Thermo Fisher Scientific) according to the manufacturer’s recommendations. The larger fragment was gel purified and used for In-Fusion cloning with a gBlock (Integrated DNA Technologies) encoding an overlapping overhangs of the human U6 promoter and the sgRNA scaffold, the eGFP stuffer and the missing part of the improved sgRNA scaffold. The In-Fusion cloning product was again transformed into One Shot Stbl3 Chemically Competent E.coli (Life Technologies) and the correct plasmid verified by Sanger sequencing.

### Cell lines and cell culture

Wild-type HAP1 C631 cells were ordered from Horizon Discovery and maintained at 37°C, 5% CO_2_ in IMDM (Life Technologies) supplemented with 10% FCS. For stable Cas9 expression, the parental cell line was transduced at an MOI of ∼0.5 with lentivirus generated using the Lenti Cas9-2A-Blast plasmid (kind gift of the Moffat lab, Addgene #73310) and selected with 20 µg/ml blasticidin (InvivoGen) for at least one week. Out of this bulk population, single cells with a small cell size (indicative for a haploid genotype) were sorted in 96-well plates, expanded and again selected with 20 µg/ml blasticidin. Blasticidin-resistant clones were further characterized for Cas9 editing efficiency. Mycoplasma contamination was periodically assessed for all cell lines. The HAP1 cell line was authenticated using Multiplex Cell Authentication by Multiplexion (Heidelberg, Germany) as described recently (Castro et al. 2012). The SNP profile was unique.

### Sorting of haploid and diploid HAP1 populations

Enrichment of haploid and diploid HAP1 Cas9 SCC11 and SCC12 populations was achieved using Flow Cytometry as described previously (Olbrich et al. 2017). In brief, respective HAP1 cells were trypsinized and a sub-sample (∼ 8*10^5^ cells) was stained using 10 µg/ml Hoechst 33342 for 30 min at 37°C. Stained cells were analyzed by flow cytometry and DNA Hoechst intensity peaks allowed for back gating of the haploid and diploid populations of interest in the FSC and SSC, as haploid cells are smaller in size. Respective gates in the FSC/SCC were used for sorting haploid and diploid HAP1 cells from unstained samples.

### Lentivirus production and MOI determination

Low-passage (<15) HEK293T cells were seeded to reach 70-80% confluency on the day of transfection. The lentiviral packaging vector psPAX2 (kind gift from the Didier Trono lab, Addgene #12260) and the lentiviral envelope vector pMD2.G (kind gift from the Didier Trono lab, Addgene #12259) were co-transfected with the respective lentiviral expression vector (∼1:1:3 molar ratio) using Trans-IT (VWR) and OptiMEM (Gibco). Roughly 16 hours post transfection, medium was replaced by fresh culture medium and lentiviral supernatant was harvested 48 hours post transfection by filtration through a 0.45 µm PES membrane, aliquoted and stored at −80 °C until transduction. For multiplicity of infection (MOI) determination, target cells were transduced with different amounts of lentiviral supernatant in the presence of 8 µg/ml polybrene. Transduced cells were selected for 48 hours with 2 µg/ml puromycin (Biomol) starting 24 hours post transduction and the number of surviving cells was compared to a non-transduced control sample.

### Single sgRNA cloning

Single sgRNA sequences were cloned into the HD CRISPR vector as described previously (Sanjana et al. 2014). In brief, the HD CRISPR vector was sequentially digested with BfuAI (NEB) and BsrGI-HF (NEB), followed by dephosphorylation using CIP (NEB). The digested backbone was gel purified using the Machery Nagel NucleoSpin Gel and PCR Clean-up kit.

sgRNA inserts were designed as two complementary oligos encoding the sgRNA target region as well as the cloning specific overhangs and were ordered as standard desalted oligos from Eurofins Genomics. Oligos were phosphorylated and annealed using T4 PNK (NEB) in 10X T4 Ligation Buffer (NEB). Diluted oligo duplexes were ligated into the digested vector using Quick Ligase (NEB) and transformed into Stbl3 recombination-deficient bacteria. A Midi- or Maxiprep (QIAfilter plasmid kit) was performed for lentivirus production or cell transfection.

### Plasmid transfection

To address functional sgRNA expression from the HD CRISPR vector, sgRNAs targeting the essential gene RNA Polymerase ll Subunit E (POLR2E) and control sgRNAs designed to target the safe harbour locus AAVS1 were cloned into the HD CRISPR vector as described above and transfected into HAP1 Cas9 bulk, HAP1 Cas9 SCC11 and HAP1 Cas9 SCC12 cells using Fugene transfection reagent (Promega). 24 hours post transfection, successfully transfected cells were selected using 2 µg/ml puromycin (Biomol) for 48 hours. Dead cells were removed by washing with PBS and viable and still attached cells were stained using 0.5 % Crystal violet (Sigma) staining solution supplemented with 20 % methanol.

### siRNA transfection

Cells were reverse transfected with 20 nM of the indicated siRNAs using Lipofectamine RNAiMAX Transfection Reagent (Thermo Fisher Scientific) and OptiMEM (Gibco). Cell viability was analyzed 48 hours post transfection.

### Flow cytometry analysis

SgRNA sequences targeting surface markers were cloned into the HD CRISPR vector as described above. Lentivirus was generated from all expression vectors. 1*10^5^ HAP1 Cas9 bulk, HAP1 Cas 9 SCC11 and HAP1 Cas9 SCC12 cells supplemented with 8 µg/ml polybrene were reversely transduced using 5 µl viral supernatant and successfully transduced cells were selected with 2 µg/ml puromycin (Biomol) 24 hours post transduction. 5 days after transduction, cells were harvested and the respective surface markers stained with an APC anti-human CD46 antibody (Biozol Diagnostica) or an APC anti-human CD81 antibody (Biozol Diagnostica). The percentage of CD46 and CD81 knockout was determined as the percentage of APC negative single cells in three independent replicates.

### Library cloning

Oligos encoding the HD CRISPR Library A and B sgRNA sequences and flanking regions were ordered as an oligo pool from Twist Biosciences. Oligos were amplified using the KAPA HiFi HotStart ReadyMix (Roche) using flanking primers in 12 PCR cycles. The resulting PCR product was purified using the NucleoSpin Gel and PCR Clean-up kit (Machery Nagel) and the correct fragment size was confirmed using a High Sensitivity Bioanalyzer DNA Kit (Agilent). For cloning, the HD CRISPR vector was digested with BfuAI (NEB) and BsrGI-HF (NEB) overnight and dephosphorylated using CIP (NEB). The resulting linearized ∼7000 bp vector missing the eGFP stuffer was gel purified again using the NucleoSpin Gel and PCR Clean-up kit (Machery Nagel) and used for library cloning in a one step digestion-ligation-reaction (Engler et al. 2008). The cloning product was purified by isopropanol precipitation and transformed into Endura ElectroCompetent Cells (Biocat), individual transformation reactions were pooled and plated on freshly prepared LB-carbenicillin plates. A 1:10 000 and 1:100 000 fold dilution was used to estimate the total number of colonies and thus the library coverage during transformation, which was aimed for to be at least 250-fold. Bacteria were incubated at 30 °C for maximal 16 h and harvested by scraping. The resulting plasmid pool was purified with the QIAfilter Plasmid Mega Kit (Qiagen). Representation and distribution of sgRNAs was analyzed by next generation sequencing.

### Analysis of non-digested backbone contaminations in plasmid preparations

To address cloning efficiency and background of remaining non-digested vector in HD CRISPR Library preparations, plasmid purifications were transfected into adherent HAP1 C631 cells using Fugene (Promega). Remaining eGFP stuffer in the non-digested vector will lead to GFP expression upon transfection and thus serves as an indicator for pooled cloning efficiency. To assess the resulting fraction of GFP positive cells upon transfection of a known proportion of undigested vector backbone, equally concentrated plasmid purifications of the non-digested HD CRISPR vector and a plasmid purification of the HD CRISPR vector expressing an AAVS1-targeting sgRNA were mixed at the indicated ratios. Successfully transfected cells were selected 24 hours post transfection using 2 µg/ml puromycin (Biomol) for 48 hours. Subsequently, cells were either imaged on an IN Cell Analyzer 6000 to address GFP expression or harvested, washed with PBS and GFP expression was analyzed by flow cytometry.

### Nicoletti ploidy stain

For DNA content analysis, HAP1 cells were harvested by trypsination and 5*10^5^ cells were washed with cold PBS by centrifugation. Cells were subsequently resuspended in 400 µl Nicoletti buffer (0.1% sodium citrate, 0,1% Triton X-100, 0.5 units/ml RNase A, 50 µg/ml propidium iodide (PI)) and incubated for 2 h at 4°C under rotation. Haploid, diploid and a 1:1 mixture of haploid and diploid HAP1 cells were prepared as controls. DNA content analysis was performed by measuring PI intensity using flow cytometry. Controls were used to set PI intensity peaks at 50 K for G1-phase haploid cells and 100 K for G1-phase diploid cells.

### Pooled CRISPR depletion screen

Prior to screening, each cell line was re-selected with 20 µg/ml blasticidin (InvivoGen) for at least one week. A total of 225 *10^6^ HAP1 Cas9 bulk, HAP1 Cas9 SCC11 and HAP1 Cas9 SCC12 cells were transduced with the lentiviral HD CRISPR Library A or B to achieve an initial library coverage of at least 300-500 fold upon infection at an MOI of ∼ 0.1-0.3, including non-transduced control flasks and accounting for variation in transduction efficiency on a large scale. 24 hours post transduction, successfully transduced cells were selected with 2 µg/ml puromycin (Biomol). One plate was cultured in the absence of puromycin for MOI determination. 48 hours post selection, cells were harvested and the MOI was determined by calculating the ratio of living cells in the presence and absence of puromycin selection. Cells were re-seeded at a 500-fold library coverage in two independent replicates for each cell line and cultured for 14 days with regular cell splitting every two to three days. A 500-fold library coverage was maintained throughout the screen for each replicate and 45 *10^6^ cells representing a ∼500-fold library coverage plus some extra cells accounting for counting and seeding errors were collected at every passage. Genomic DNA was isolated from the final passage for genomic DNA extraction and sequencing.

### Genomic DNA extraction, library preparation and sequencing

Genomic DNA was isolated from frozen cell pellets using the QIAAmp DNA Blood and Tissue Maxi Kit (Qiagen) and purified by ethanol precipitation. The sgRNA spanning region was amplified from purified genomic DNA at a 100-fold library representation. Illumina adapters and indices were added in the same one-step PCR reaction using the KAPA HiFi HotStart ReadyMix (Roche). The PCR product was purified using the QIAQuick PCR Purification Kit (Qiagen) and eluted on MiniElute columns (Qiagen) upon library preparation from a plasmid pool or further gel purified to remove genomic DNA contamination using the NucleoSpin Gel and PCR Clean-up kit (Machery Nagel) in case of library preparation for screening samples. The correct fragment size was confirmed with a High Sensitivity Bioanalyzer DNA Kit (Agilent). Sequencing was performed on an Illumina NextSeq 550 system with a High-Output Kit (75 cycles) (Illumina). Low heterogeneity was either addressed by using a custom sequencing primer binding immediately upstream of the sgRNA targeting sequence or by using the standard Illumina sequencing primer and adding 20% PhiX.

### Calculation of sgRNA sequence scores

For the calculation of Ruleset 2 scores (Doench et al. 2016) flanking regions for each sgRNA were retrieved using the Bioconductor BSgenome package (human genome version hg38) (Huber et al. 2015). Ruleset 2 scores were then calculated using the previously published software (Doench et al. 2016). Similarly, DeepHF scores were computed using published software (Wang et al. 2019) according to the instructions on the corresponding GitHub page (https://github.com/izhangcd/DeepHF). Finally, optimized sgRNA scores according to Hart *et al.* were calculated based on the position weight matrix published with the corresponding manuscript (Hart et al. 2017).

### Initial data processing and analysis

The MAGeCK software version 0.5.7 (Li et al. 2014) was used to quantify sgRNA abundance from sequencing data. A pseudo count was added for each sgRNA in each sample. To adjust for differences in sequencing depth, samples were then normalized by dividing sgRNA counts to the median count of the targeting controls. sgRNA depletion phenotypes were quantified as log_2_ fold changes were calculated for each sgRNA as 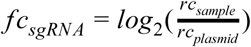 where *rc*_*sample*_ are normalized read counts of samples after 14 days of selection and *rc*_*plasmid*_ are normalized counts of the plasmid library. Reproducibility between replicates was assessed using Pearson and Spearman correlation coefficients. To combine the screens in sub-libraries A and B into a combined library dataset, raw counts for each sample were first normalized to the same library depth through division by the median count in each sample and multiplication by the median count across all samples. Normalized counts for screens in sub-libraries A and B were then combined into a combined library count file. Normalized counts were further rounded to the closest count. To determine knockout phenotypes of house-keeping (core essential) genes compared to nonessential genes previously reported gold standard lists of core (CEGv2) and nonessential (NEGv1) genes were used (Hart et al. 2014, 2017).

### Classification of essential genes

To evaluate screen performance, BAGEL v0.91 (Hart and Moffat 2016) was used to classify genes as essential and nonessential (Zhan and Boutros 2016). Specifically, a gene was classified as essential if the Bayes Factor (BF) determined by BAGEL was greater than 6 (similar cutoffs were chosen as in (Hart et al. 2017)). Precision-recall curves and statistics to quantify how well core and nonessential reference genes could be separated based on each screen’s depletion phenotypes were determined using the ROCR R package (Sing et al. 2005). To compare essential gene detection power of individual sub-libraries compared to the combined library, the MAGeCK RRA (Li et al. 2014) and gscreend (Imkeller et al. 2019) algorithms were used in addition to BAGEL to determine essential genes at 5% FDR. MAGeCK RRA was used with default paramters. For gscreend analysis the number of permutations used to calculate the ρ_0_ parameter for gene ranking was set to 10,000.

### Analysis of cutting and non-cutting sgRNAs

A Gaussian mixture model with 4 components was fit to the fold change distributions of all targeting (n=270) and non-targeting control (n=600) for each screen separately using an expectation maximization algorithm implemented in the R package ‘mixtools’ (Benaglia et al. 2009). Here, two components were used to capture the fold change distributions of targeting and non-targeting controls (henceforth referred to as components 1 and 2, respectively) and two additional mixture components were used to capture moderate and several viability phenotypes of toxic controls (components 3 and 4; Supplementary Figure 8B). The resulting models were used to classify each additional sgRNA in the library as follows: each sgRNA was mapped to the control with the most similar phenotype. If that control was assigned to components 3 or 4 with a probability of at least 80% the sgRNA was assumed to have a target-dependent viability phenotype. If the associated control was assigned to component 1 or 2 with a probability of at least 80% an sgRNA was considered ‘likely cutting’ or ‘likley not cutting’, respectively. If a control could not confidently be assigned to any of the mixture components (probability for all components < 80%) then all sgRNAs with similar phenotypes were labeled ‘undetermined’.

### Influence of library coverage on essential gene detection

To investigate how library coverage affects the number of essential genes that can be detected, 2 to 8 sgRNAs per gene were sampled from the combined HD CRISPR Library. Five independent samplings were performed for each library coverage. BAGEL v0.91 (Hart and Moffat 2016) and MAGeCK RRA v0.5.7 (Li et al. 2014) were then used to classify essential genes at BF > 6 (for BAGEL) and FDR < 5% (for MAGeCK) in both the bulk population and the single cell clones. CEGv2 and NEG1 core- and nonessential reference gene sets were used for essential gene detection with BAGEL.

### Data and software access

All raw sequencing data generated in this study have been submitted to the European Nucleotide Archive (ENA; https://www.ebi.ac.uk/ena) under the accession number PRJEB35190. Documented computer code to reproduce all figures presented in this study is available through GitHub at https://github.com/boutroslab/Supplemental-Material/tree/master/Henkel%26Rauscher_2019.

## Acknowledgements

We thank Fillip Port for help with the design of the cloning strategy and Josephine Bageritz for help with FACS analysis. We further thank the members of the Boutros laboratory for hands-on help of large-scale library cloning experiments and helpful discussions and feedback, especially Fillip Port and Katharina Imkeller for helpful comments on the manuscript. We are grateful for the excellent support of the DKFZ Genomics and Proteomics Sequencing core facility in sequencing screening samples, especially to N. Diessl, as well as the DKFZ Flow Cytometry core facility. We further thank the ZMBH FACS core facility (M. Langlotz) for help with sorting single cell clones and the EMBL Genomics Core Facility, especially V. Benes, for excellent sequencing support. Furthermore, we thank Matthias Hinterndorfer and Johannes Zuber for providing CD46 sgRNA sequences and the scientific community for sharing CRISPR screening data. B.R. was supported by the BMBF-funded Heidelberg Center for Human Bioinformatics (HD-HuB) within the German Network for Bioinformatics Infrastructure (de.NBI) (Grant #031A537A). The project has been supported in part by an NCT Basic Science Grant. Work in the Boutros laboratory is supported in part by an ERC Advanced Grant.

---

**Supplementary Figure 1:**
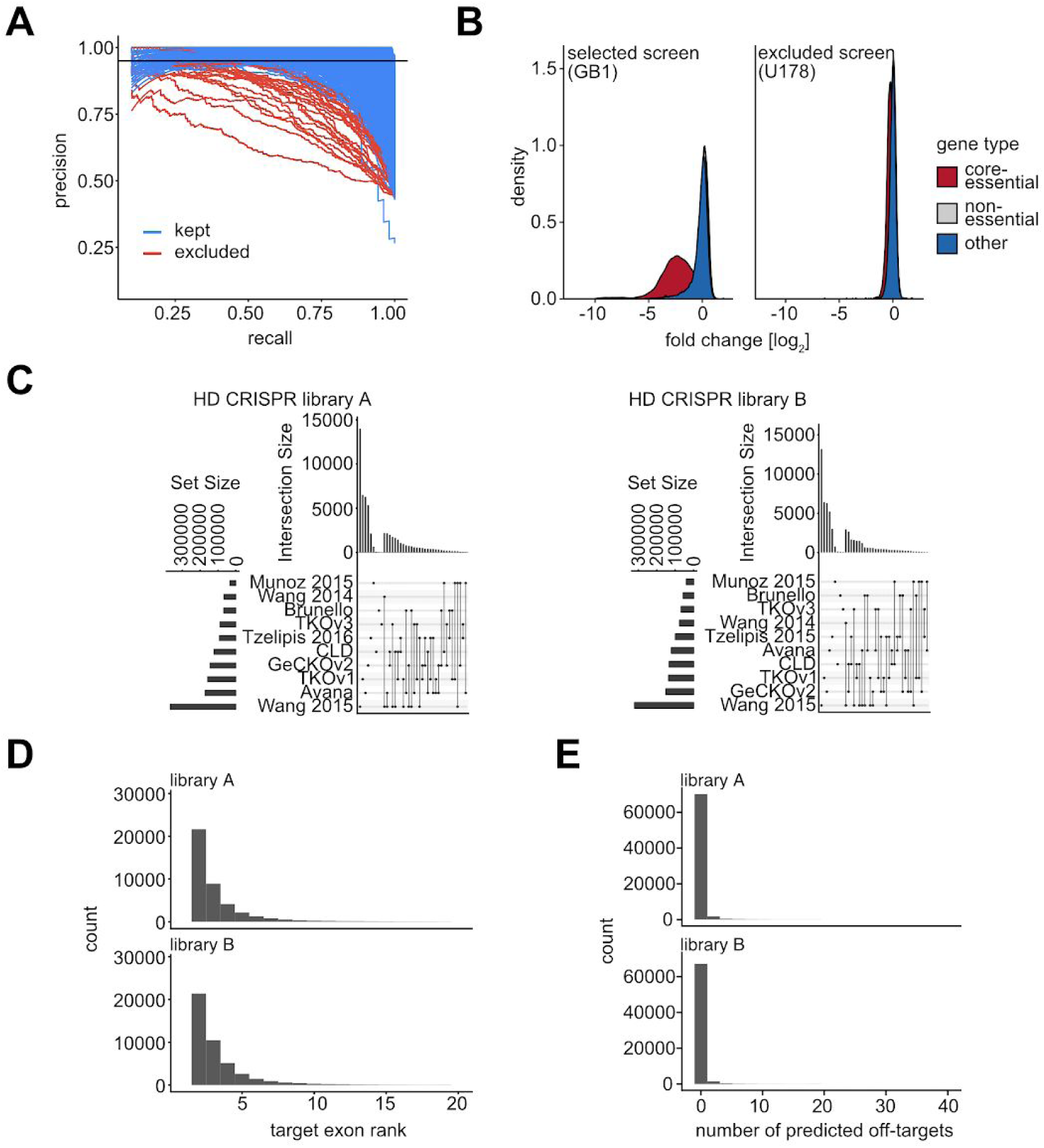
HD CRISPR Library composition. (A) Precision recall curves for differentiating reference core and nonessential genes based on BAGEL Bayes factors determined for published fitness screens in GenomeCRISPR. Blue curves indicate screens with an area under the curve (AUC) greater than 0.9. Red screens with an AUC of less than 0.9 were excluded for HD CRISPR sgRNA design. (B) Log2 fold change distributions of core essential (red) and nonessential (blue) reference genes for a high quality (left) and a low quality (right) example screen. (C) Library composition of the HD CRISPR sub-libraries A and B. Horizontal bars on the left indicate the number of designs used from different previously published libraries. The panel on the bottom right shows combinations of libraries, that include designs selected for the HD CRISPR Library. The bars above this panel quantify the number of selected sgRNAs for each of these combinations. (D) Distribution of exon ranks targeted by the sgRNAs in the HD CRISPR Library. (E) Distribution of the predicted off-target counts (see Materials & Methods) for sgRNAs in the HD CRISPR Library.

**Supplementary Figure 2:**
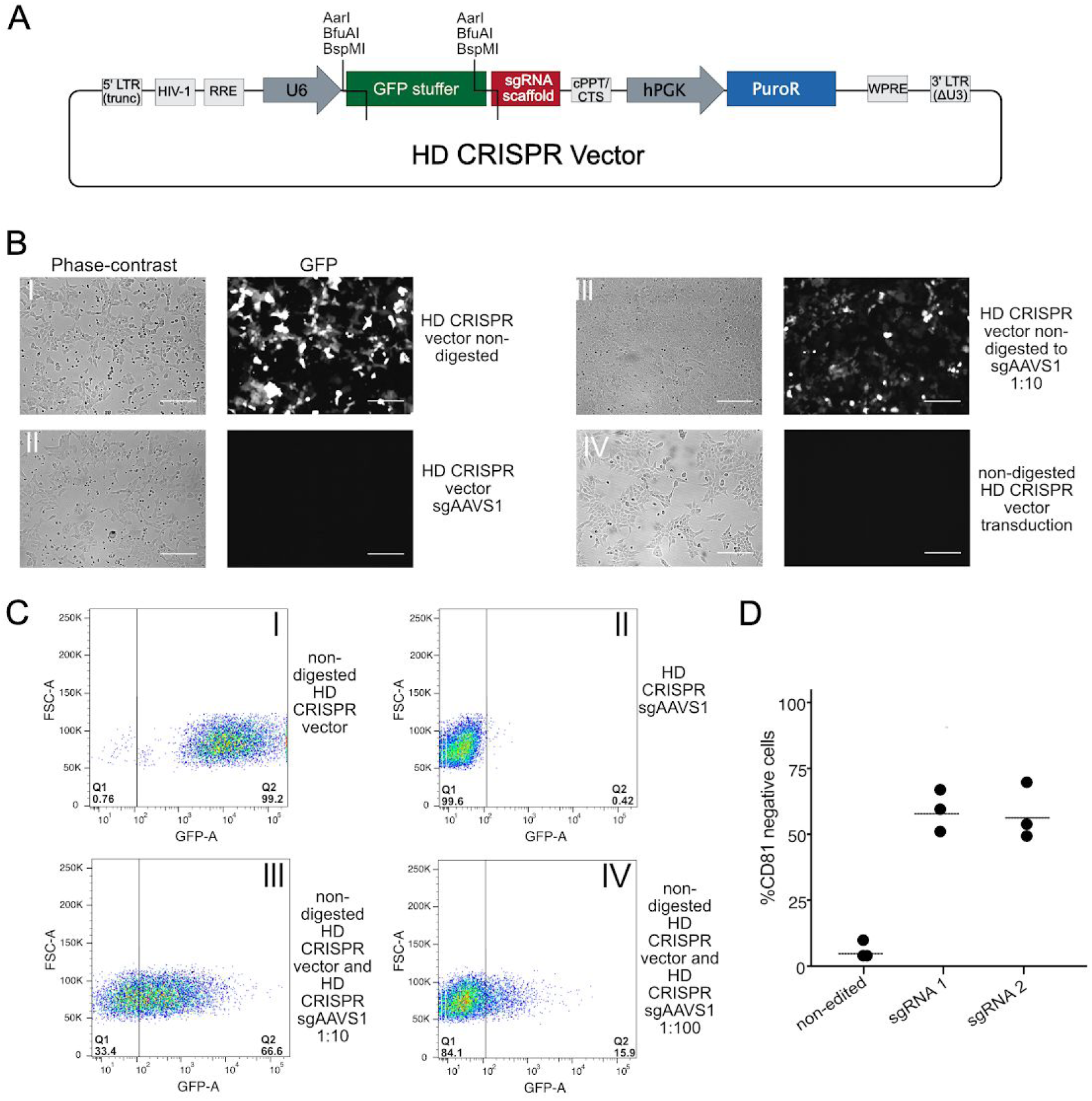
Features and performance of the HD CRISPR vector. (A) Composition of the lentiviral HD CRISPR sgRNA expression vector. (B) sgRNA cloning efficiency can be addressed upon transfection of the HD CRISPR vector, since residual GFP stuffer in non-digested vector backbone leads to GFP expression (B.l). Complete removal as achieved when cloning single sgRNAs abolishes GFP expression (B.ll), while remaining stuffer in 10% of the plasmid pool still leads to a substantial amount of GFP positive cells (B.lll). Transduction of the non-digested vector still containing the GFP-stuffer does not result in GFP-expressing cells (B.IV). Representative images from two independent experiments are shown. Scale bar = 100µM (C) FACS analysis of GFP expression upon transfection of the non-digested HD CRISPR vector (l) or the HD CRISPR vector expressing an sgRNA (ll). A mixture of GFP positive and negative cells can be observed upon transfection of a mixture of stuffer and sgRNA-containing vector (lll and IV). (D) Editing efficiency was furthermore assessed upon transduction of HAP1 Cas9 cells with the HD CRISPR vector expressing sgRNAs targeting the surface proteins CD81, followed by FACS staining of residual CD81 protein to address knockout efficiency. Antibody staining of the non-edited cell line was used as a control. Shown are representative images from three independent experiments except for the 1:100 dilution, where only two measurements were taken.

**Supplementary Figure 3:**
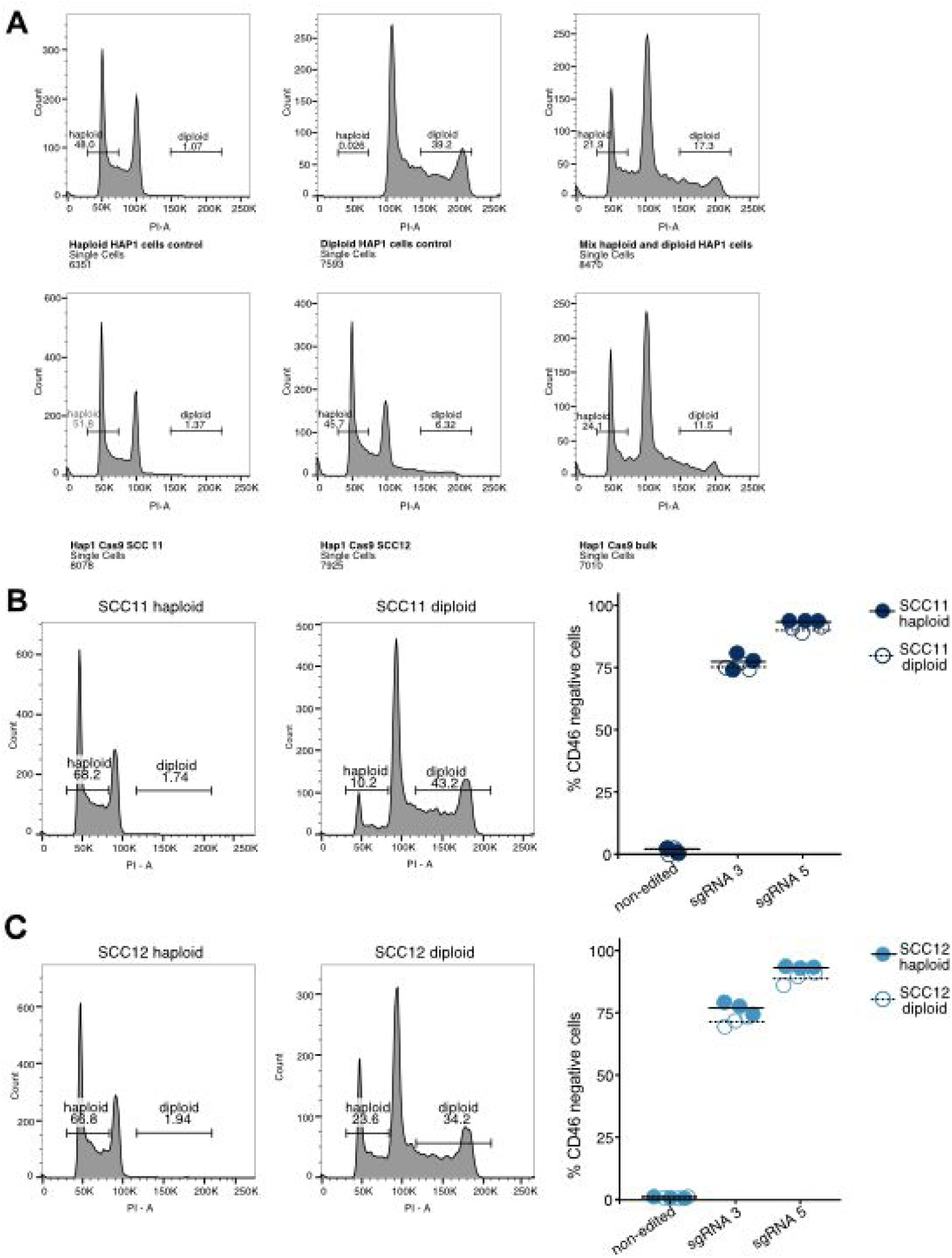
DNA content analysis to determine ploidy of various HAP1 Cas9 populations. (A) HAP1 Cas9 bulk and HAP1 Cas9 SCC11 and SCC12 cells were stained for DNA content using Nicoletti buffer and FACS analysis. The percentage of G1 haploid cells and G2 diploid cells are indicated for each cell population. (B-C) Enriched haploid and diploid populations of the HAP1 Cas9 SCC11 (B) and Cas9 SCC12 (C) cell lines were obtained by FACS sorting. Subsequently, haploid and diploid populations were independently transduced with the HD CRISPR vector expressing sgRNAs targeting the surface marker *CD46* and editing efficiency was directly compared in the haploid and diploid population of the same cell line. Non-edited samples of the respective cell lines served as a control.

**Supplementary Figure 4:**
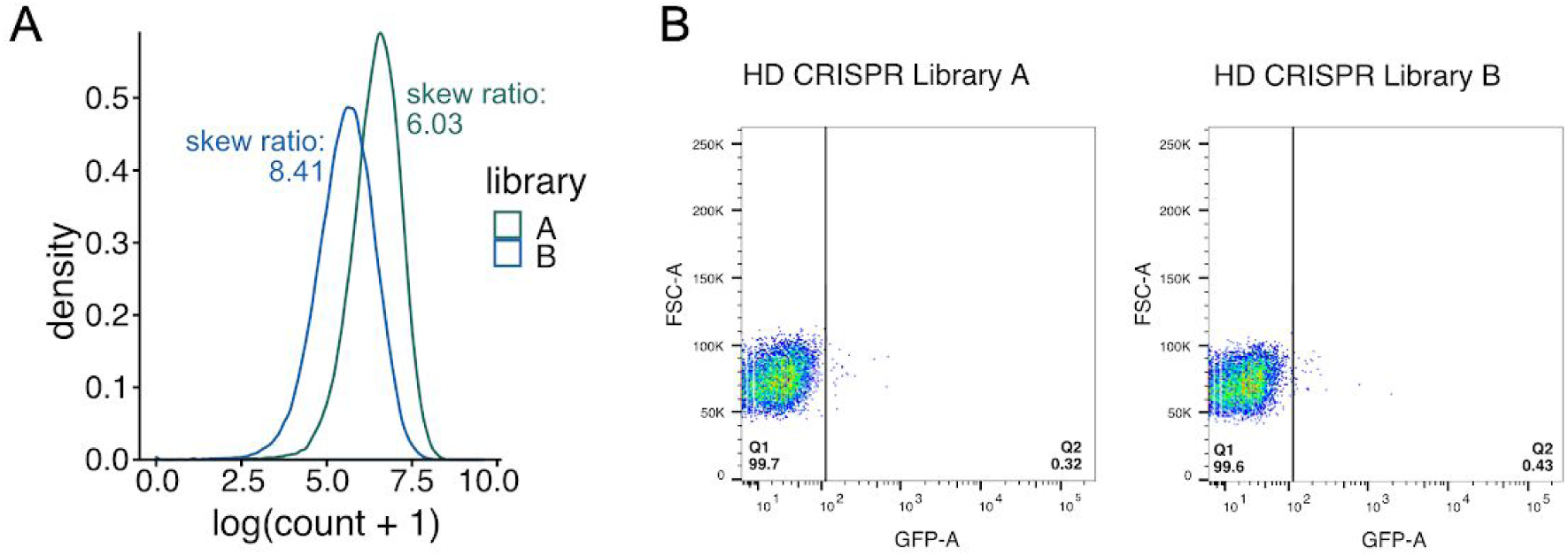
Cloning quality control of the HD CRISPR Library. (A) Distribution of sgRNA read counts for the HD CRISPR plasmid library preparations. Skew ratios were determined as the quotient of the top 10 quantile divided by the bottom 10 quantile. (B) FACS analysis of GFP expression upon transfection of the HD CRISPR Library A and B plasmid pools to address the presence of remaining GFP stuffer.

**Supplementary Figure 5:**
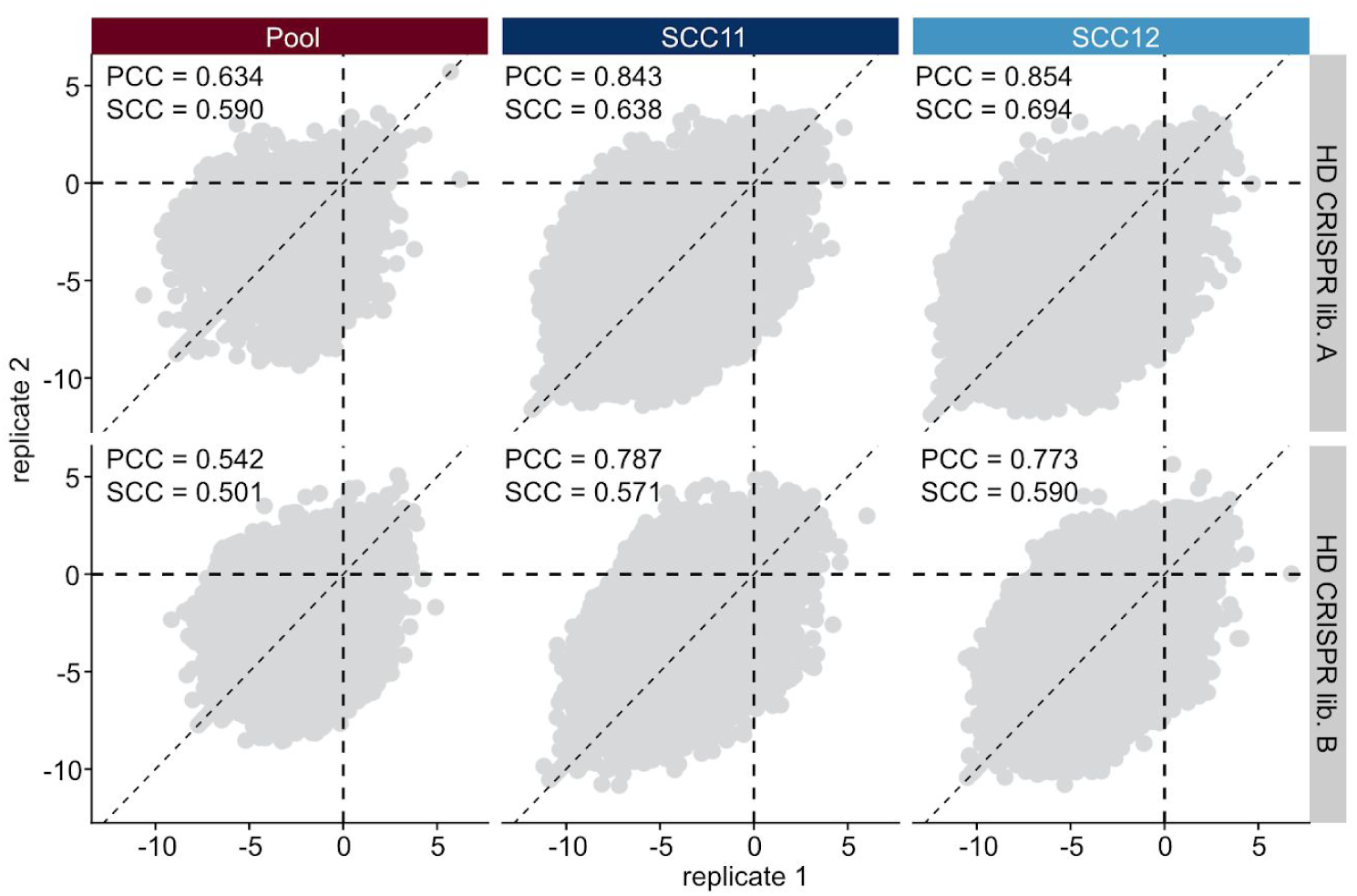
Reproducibility of negative selection screens with the HD CRISPR Library. Scatter plots showing the reproducibility of sgRNA phenotypes across biological replicates in screens with the HD CRISPR Library. Each column includes screens performed in a bulk cell population (left) or in selected single cell clones with high Cas9 activity (middle and right). The top and bottom rows include screens with the HD CRISPR sub-libraries A and B, respectively. PCC = Pearson correlation coefficient, SCC = Spearman correlation coefficient.

**Supplementary Figure 6:**
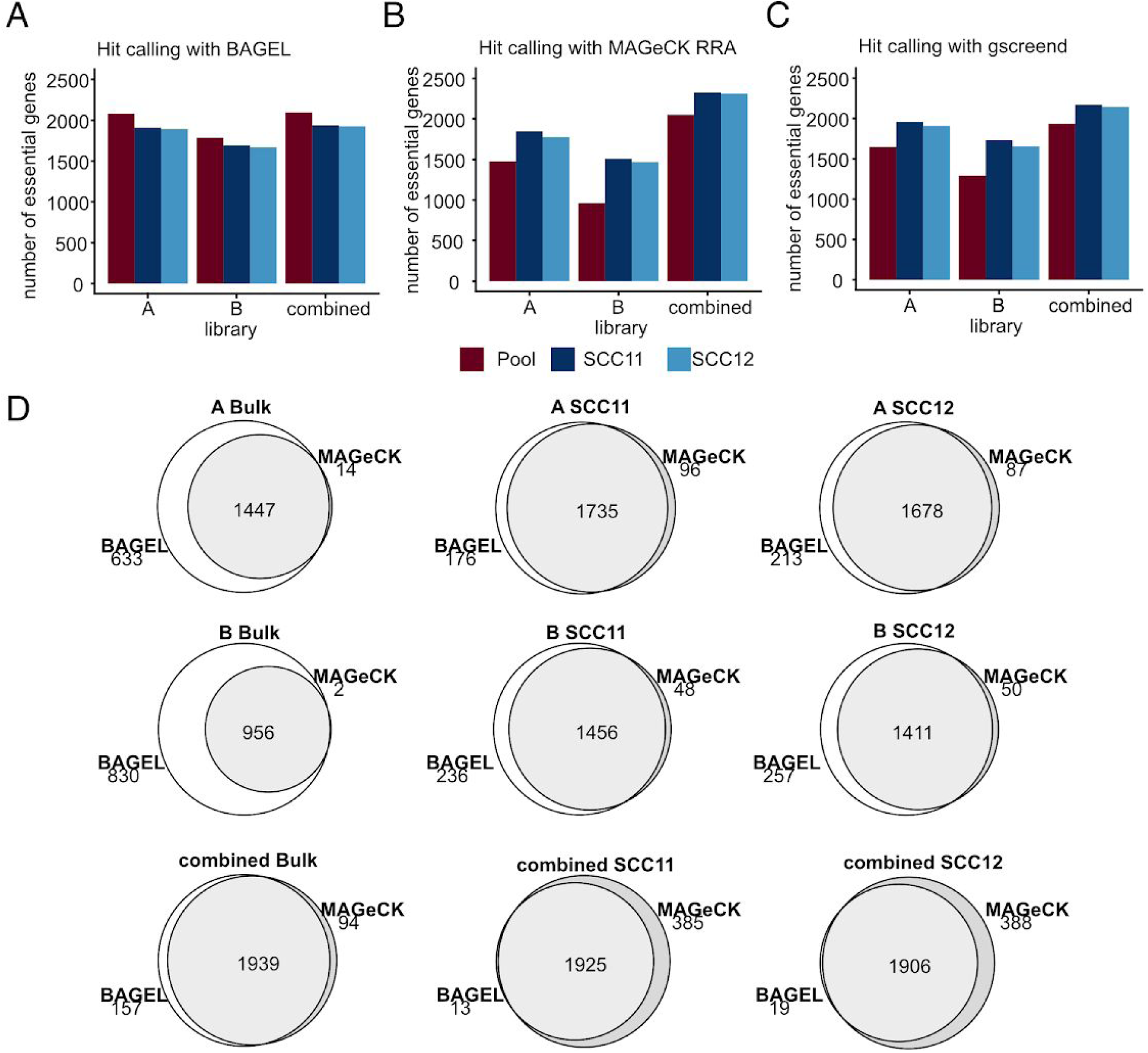
Hit detection in screens with the HD CRISPR Library. (A) Number of hits determined using BAGEL (Hart and Moffat 2016) at a strict Bayes factor cutoff (BF > 6) in different screens conducted with the HD CRISPR Library. (B) Number of essential genes determined using MAGeCK RRA (Li et al. 2014) at 5% FDR in different screens conducted with the HD CRISPR Library. (C) Number of essential genes determined using gscreend (Imkeller et al. 2019) at 5% FDR in different screens conducted with the HD CRISPR Library. (D) Venn diagrams show overlap between essential genes determined using either BAGEL or MAGeCK RRA for each screen.

**Supplementary Figure 7:**
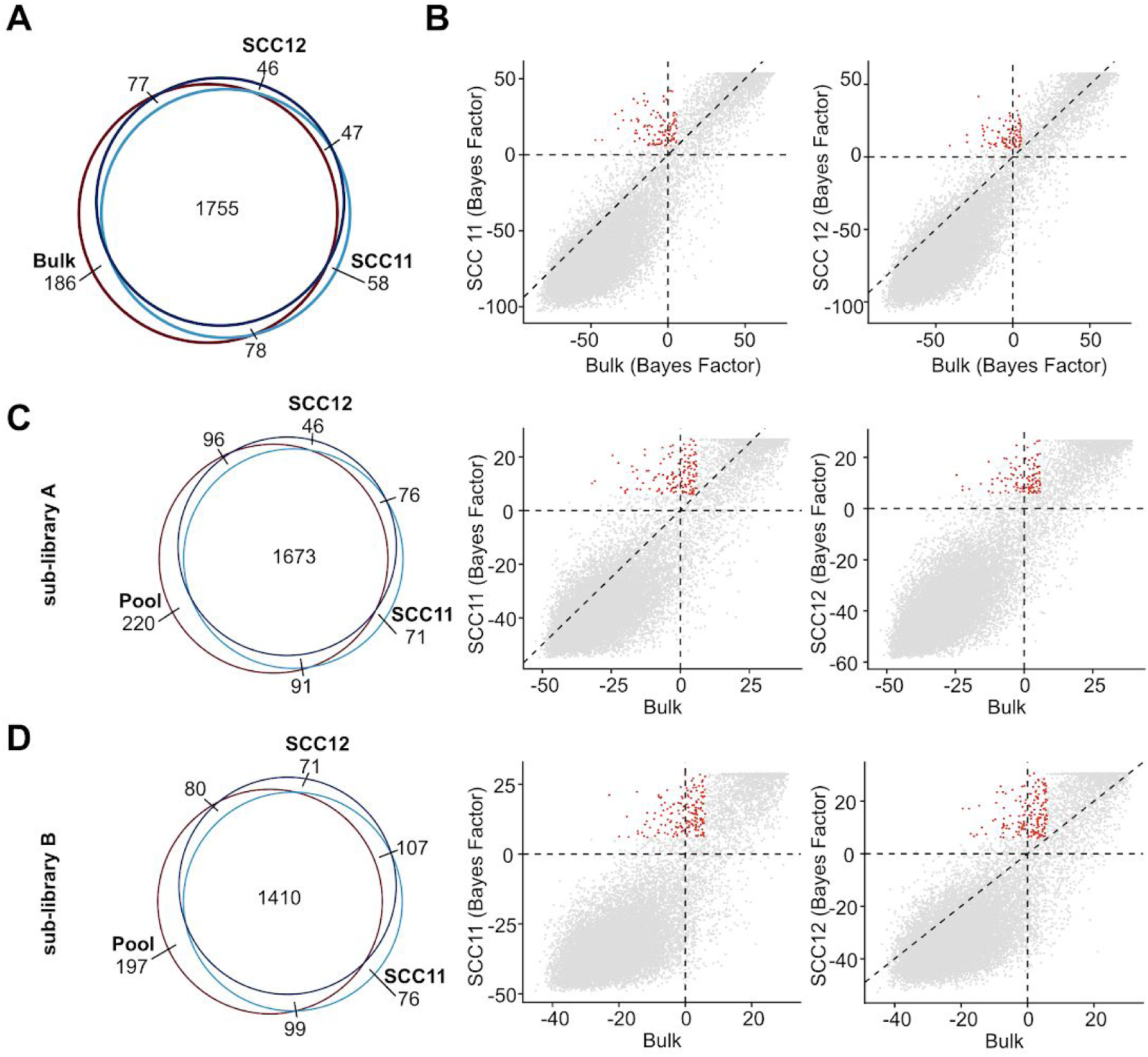
Essential genes are highly consistent between HAP1 Cas9 bulk population and single cell clones. A) Venn diagram showing essential gene overlap between a HAP1 bulk Cas9 population and two single cell clones that were selected for high Cas9 activity. Gene essentiality was determined using BAGEL with a Bayes Factor cutoff of 6 (see (Hart et al. 2017)). The combined HD CRISPR Library with 8 sgRNAs per gene was used for essential gene inference. (B) Quantitative comparison of BAGEL Bayes Factors for each gene between the HAP1 bulk Cas9 population and selected single cell clones SCC11 and SCC12. Each dot represents a gene in the HD CRISPR Library. Red dots indicate essential genes that are private to a single cell clone. The dashed diagonal is the identity line. (C) Venn diagram (left) and scatter plots (middle and right) showing essential gene overlap between a HAP1 bulk Cas9 population and two single cell clones in screens using the HD CRISPR sub-library A. (D) Venn diagram (left) and scatter plots (middle and right) showing essential gene overlap between a HAP1 bulk Cas9 population and two single cell clones in screens using the HD CRISPR sub-library A.

**Supplementary Figure 8:**
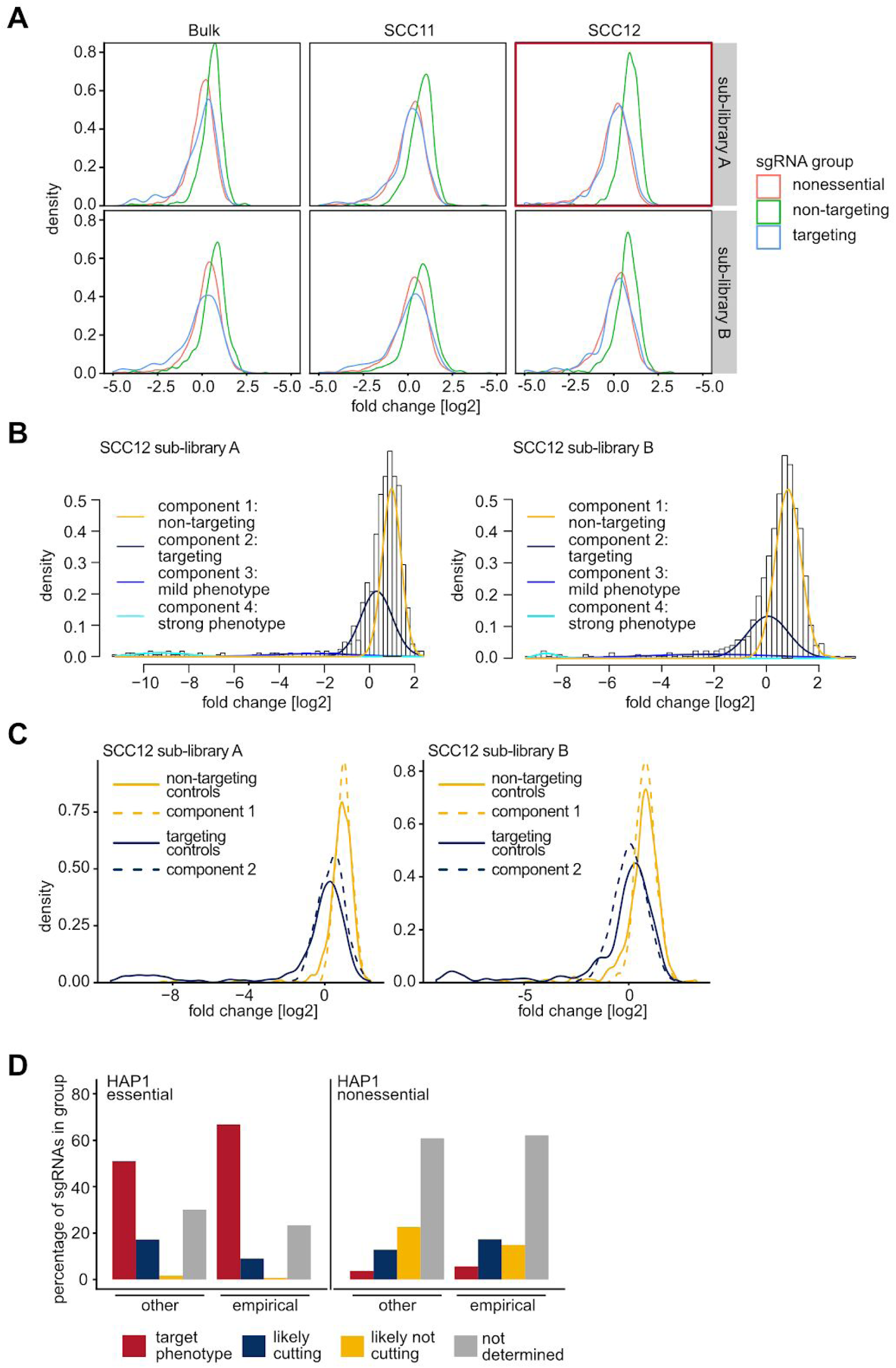
Prediction of sgRNA DNA cutting activity based on control phenotypes. (A) Log2 fold change phenotype distributions for sgRNAs targeting nonessential genes (red) as well as targeting (blue) and non-targeting control sgRNAs (green) across different screens conducted with the HD CRISPR Library. The screen with the HD CRISPR Library A in HAP1 single cell clone SCC12, which was used for subsequent analyses, is highlighted in red. (B) Fit of a Gaussian mixture model with 4 components for screens in SCC12. Components 1 (yellow) and 2 (blue) represent non-targeting and targeting sgRNAs, respectively. Components 3 and 4 capture the phenotypes of sgRNAs with moderate and severe viability phenotypes. (C) Comparison of true fold change distributions of targeting and non-targeting sgRNAs (solid line) to the distributions estimated by the mixture model components (dashed lines) for both HD CRISPR libraries A and B. (D) Number of sgRNAs associated with each phenotype group targeting essential genes according to MAGeCK analysis. For this representation components 3 and 4 are combined in the red group ‘target phenotype’. sgRNAs are stratified based on their design: ‘empirical essential’ sgRNAs target context-specific essential genes and were selected for the HD CRISPR Library based on their previous on-target phenotypes. ‘Empirical nonessential’ sgRNAs are part of previously published libraries and target broadly nonessential genes. They were selected based on their lack of outlier phenotypes. *De novo* sgRNAs were designed using the software cld (Heigwer et al. 2016). (E) Similar plot as D showing sgRNAs of the HD CRISPR library B associated with each phenotype group.

**Supplementary Figure 9:**
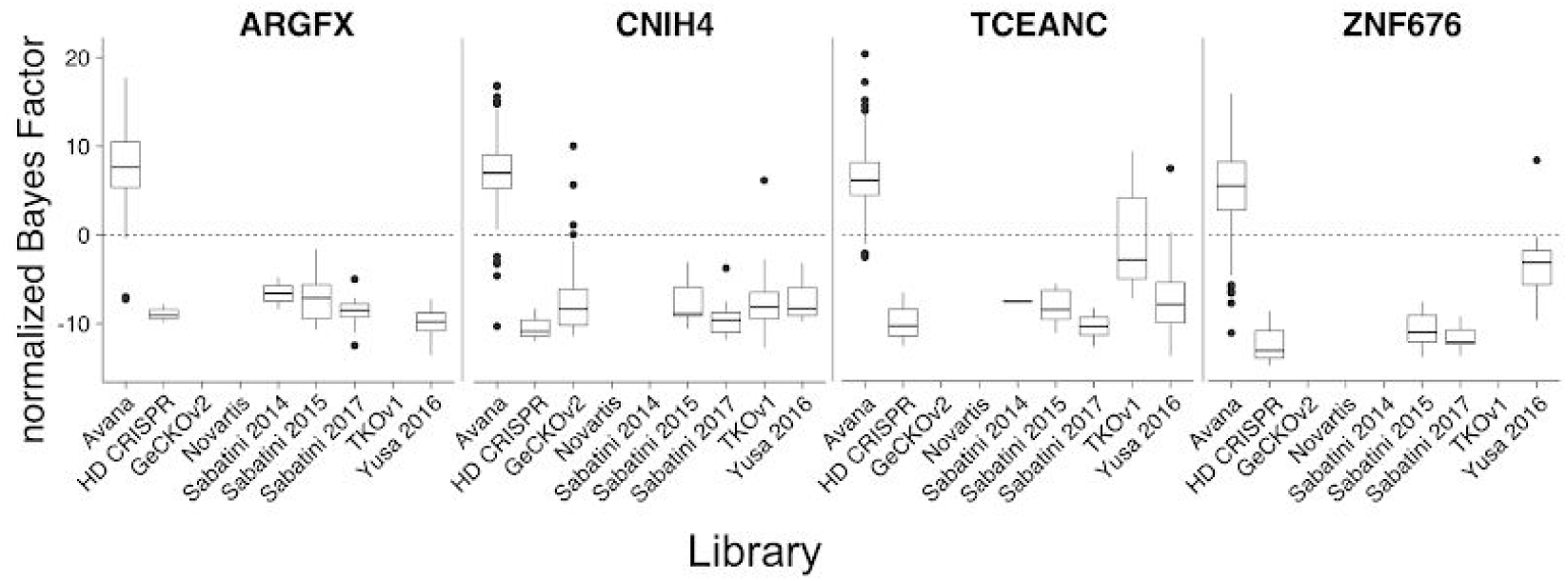
HD CRISPR Library design strategy does not enrich for sgRNAs with strong phenotypes presumably caused by off-target effects. Bayes factor analysis of selected HAP1 context-dependent nonessential genes across different screens conducted with various genome-scale CRISPR libraries in cancer cell lines.

**Supplementary Figure 10:**
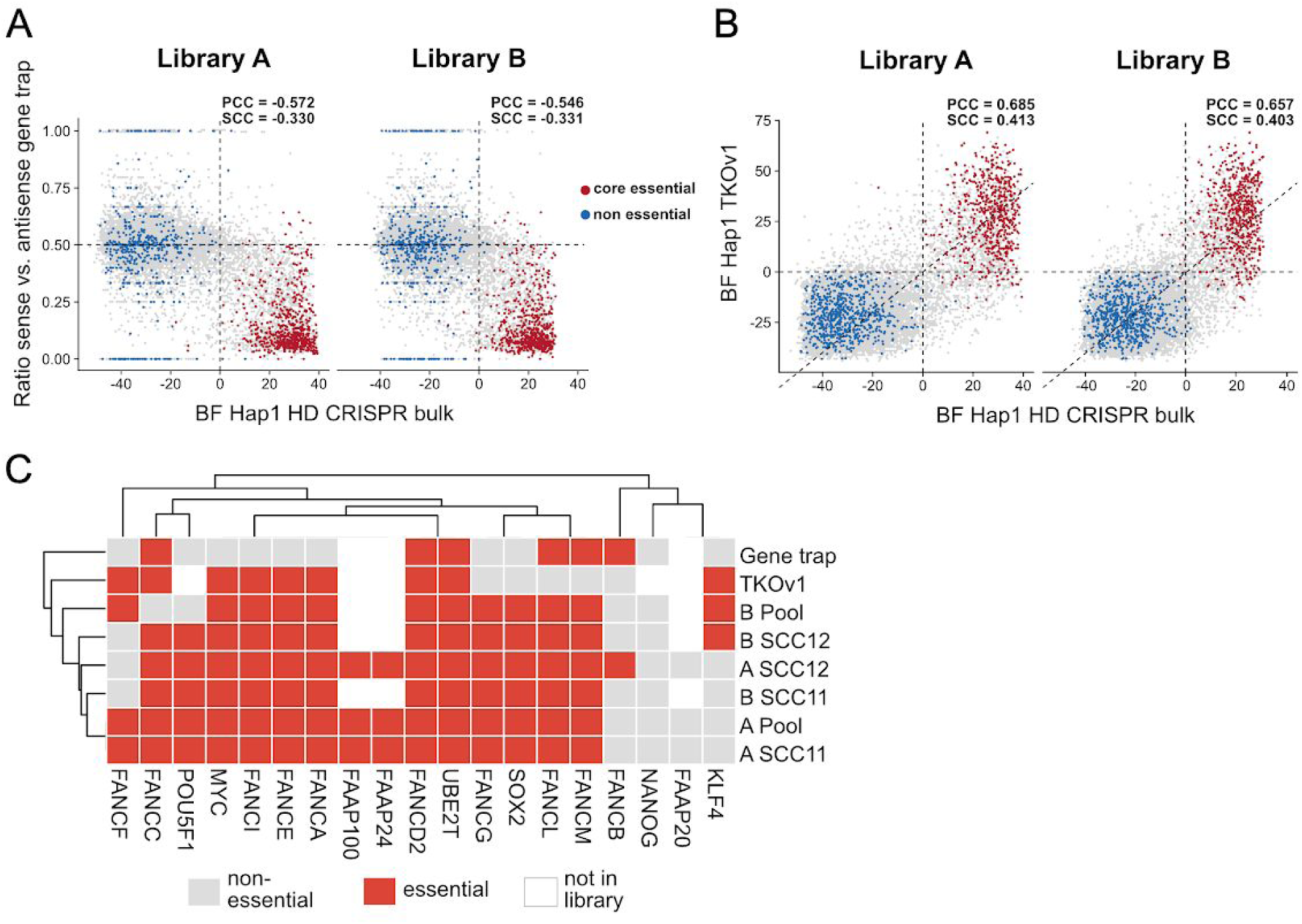
Hit calling of the HD CRISPR Library correlates with former screens conducted in HAP1 cells. (A) Hit calling of the HD CRISPR Libraries A and B in comparison with a gene trap screen conducted by Blomen et al. (2015) in HAP1 cells. (B) Hit calling of the HD CRISPR Libraries A and B in comparison with a CRISPR screen conducted in HAP1 cells by Hart et al. (2017) using the TKOv1 library. (C) Essentiality of Yamanaka factors and Fanconi anemia pathway members in previous Hap1 screens. Red boxes indicate that the gene was found essential and grey indicates non-essentiality. White boxes represent genes that are not targeted by the respective library.

